# Effects of Calreticulin Mutations on HLA Class I Expression in Myeloproliferative Neoplasms

**DOI:** 10.1101/2025.06.24.661416

**Authors:** Amanpreet Kaur, Harini Desikan, Grace Pagnucco, Malathi Kandarpa, Moshe Talpaz, Malini Raghavan

## Abstract

Calreticulin (CRT) is important for human leukocyte antigen (HLA) class I assembly. Somatic mutations of the CRT gene (*CALR*) in hematopoietic lineage cells cause myeloproliferative neoplasms (MPNs). Typically, MPN patient cells have one copy each of the wild-type and mutant *CALR* allele. We find that heterozygous knock-in of a MPN *CALR* mutation into human cell lines maintains or slightly induces surface expression of HLA class I allotypes. However, full deficiency of wild-type CRT variably reduces the surface expression of HLA class I allotypes, and MPN CRT mutants fail to restore expression for all tested allotypes. Consistent with the largely heterozygous nature of *CALR* mutations in MPN, surface HLA class I expression in platelets and monocytes from MPN patients with *CALR* mutations generally falls within the normal range, with higher average expression measured in monocytes from patients treated with interferon alpha compared with other treatments. Overall, the studies indicate that loss of HLA class I expression in cells deficient in wild-type CRT (the CRT dependency) is allele-dependent and correlates with known effects of the assembly factor tapasin. Furthermore, heterozygous knock-in of a MPN-linked *CALR* mutation has no effect on some allotypes and slightly induces HLA class I expression for CRT-dependent allotypes.

## Introduction

MPNs are a group of hematopoietic cancers characterized by abnormal expansion of myeloid lineage hematopoietic cells. *CALR* mutations have been associated with a subset of MPNs, specifically ET and myelofibrosis (MF)^1,2^. CRT is a glycoprotein chaperone primarily localized in the endoplasmic reticulum (ER) that contains multiple low-affinity calcium-binding sites near its C-terminus^3,4^. The most prevalent MPN-linked *CALR* mutations include a 52 bp deletion (Del52) and a 5 bp insertion (Ins5) within *CALR* exon 9^1,2^. Mutant CRT proteins form stable complexes with the thrombopoietin receptor (TPOR or myeloproliferative leukemia protein (MPL))^5–8^. The CRT mutants and MPL complexes are co-trafficked to the cell surface to stimulate JAK/STAT signaling and downstream protein expression that supports thrombopoietin (TPO)-independent cell proliferation and differentiation^5–7,9,10^.

CRT is important for the folding and assembly of major histocompatibility complex (MHC) class I proteins^11–13^ (reviewed in^14^). MHC class I molecules are cell surface glycoproteins expressed on all nucleated cells and platelets that present antigenic peptides to CD8^+^ T cells and mediate the activation of CD8^+^ T cells and immune responses against cancers and viral infections. CRT is a component of a multiprotein entity called the peptide loading complex (PLC) that comprises nascent MHC class I molecules bound to CRT, the oxidoreductase ERp57, and the MHC class I-specific assembly factors tapasin and the transporter associated with antigen presentation (TAP)^14–17^. The human MHC (human leukocyte antigen; HLA) class I proteins are highly polymorphic, with thousands of allelic variants prevalent in the human population^18^. These polymorphisms not only define distinct peptide-binding specificities of different allotypes^19–22^, but also induce different dependencies of HLA class I allotypes on components of the PLC for their assembly and cell surface expression^23–27^. In HIV-infected patients, the variable tapasin dependencies of HLA class I allotypes affect progression to AIDS via an influence on the breadth of CD8^+^ T cell responses^27^. TAP dependencies are also somewhat variable, and some HLA class I allotypes are more readily detectable in the presence of TAP inhibitors or TAP deficiency^26,28,29^.

In murine fibroblasts, CRT deficiency reduced MHC class I assembly and cell surface expression, which, in turn, compromised antigen presentation to CD8^+^ T cells^11^. Additionally, in HEK293T cells, CRT deficiency induced a reduction in the overall surface HLA class I expression, which was not restored by the expression of frameshift MPN-linked CRT mutants^30^. However, whether there are allele-specific differences in the effects of CRT deficiency or mutation on HLA class I expression is unknown. Furthermore, most MPN patients have cells with heterozygous *CALR* mutations, retaining one copy of the wild-type *CALR* allele^1,31^, but how heterozygous *CALR* mutations affect the expression of HLA class I allotypes is unknown. These gaps in knowledge were addressed in this study.

## Methods

### Blood donors and ethics

This study was approved by the University of Michigan Institutional Review Board. Patient blood samples and the related clinical information including the disease, mutation, treatment and variant allele frequencies were collected after rigorous de-identification from the Myeloproliferative disease repository (study ID: HUM0006778). Healthy donor blood samples were collected after obtaining written informed consent from donors recruited in a research study (study ID: HUM00071750) and from the University of Michigan Platelet Physiology and Pharmacology Core repository (study ID: HUM00107120).

### DNA constructs

The single guide RNA (sgRNA) targeting a sequence 5’-GGCCACAGATGTCGGGACCT-3’ located at the boundary of intron 3 and exon 4 of the *CALR* gene was used for CRT knockdown^32^. The pMSCV constructs of wild-type human *CALR* (*CRT_WT_*), *CRT_Del52_*, and *CRT_Ins5_* have been described earlier^8^. For reconstitution of CRT expression in CRT-KO cells, the sgRNA-resistant versions of *CRT_WT_*, *CRT_Del52_*, and *CRT_Ins5_* generated by site-directed mutagenesis were cloned in the pSIP-*ZsGreen* (pLVX-SFFV-IRES-Puro) vector^27,33^ that was a gift from Dr. Mary Carrington at the Frederick National Laboratory for Cancer Research. The pMSCV neo constructs used for expression of HLA class I in K562 cells have been described earlier^25^. For some HLA class I allotypes, the genes were subcloned into the pSIP-*ZsGreen* vector using primers given in supplementary table 3.

### CRISPR/Cas9-based CRT knockout and knock-in

The sgRNA targeting *CALR* was cloned in the pLentiCRISPRv2-BLAST vector and was used for knockdown of CRT expression in K562, B-Lymphoblastoid Cell lines (B-LCLs), and THP-1 cells. The CRISPR-based knock-in approach described earlier^34^ was used to generate cell lines with heterozygous knock-in of the Del52 mutation at the endogenous *CALR* locus. The detailed procedures are given in the Supplementary Information.

## Results

### Calreticulin haploinsufficiency induces or maintains surface HLA class I expression for multiple allotypes

We used a previously described CRISPR knock-in approach^34^ to create cell lines with heterozygous Del52 knock-in at the endogenous *CALR* locus. HLA class I-deficient K562 lymphoblast cells and THP-1 monocytic cells were nucleofected with Cas9-ribonucleoprotein (RNP) and infected with an equal titer of adeno-associated virus (rAAV) particles packaged with either wild-type or mutant CRT-specific knock-in templates. The wild-type and Del52 knock-in templates included a GFP and BFP reporter gene, respectively, placed downstream of respective *CALR* fragments and expressed from an independent spleen focus-forming virus (SFFV) promoter. Analysis of K562 cells by flow cytometry showed the presence of ∼2-3% single positive (GFP^+^ or BFP^+^) cells while the percentage of GFP^+^BFP^+^ cells was small (<1%) (**Figure 1A**). All four cell subsets (GFP^+^, BFP^+^, GFP^+^BFP^+^ and GFP^-^BFP^-^) were sorted. Immunoblots from the sorted cells confirmed the expression of both wild-type CRT and mutant CRT proteins in the K562 BFP^+^ and GFP^+^BFP^+^ cells (**Figure 1B**, lanes 4-6) and THP-1 BFP^+^ cells (**Supplementary Figure 1A**). Wild-type CRT expression from the knock-in allele in the GFP^+^ or GFP^+^BFP^+^ cells was higher than the endogenous CRT levels in the GFP^-^BFP^-^ and BFP^+^ cells (**Figure 1B**, lanes 7-12 *vs.* lanes 1-6). This is likely caused by the SV40 polyA sequence placed at the 3’ end of the *CALR* knock-in fragment that stabilizes the *CALR* mRNA expressed from the knock-in allele.

**Figure 1:**
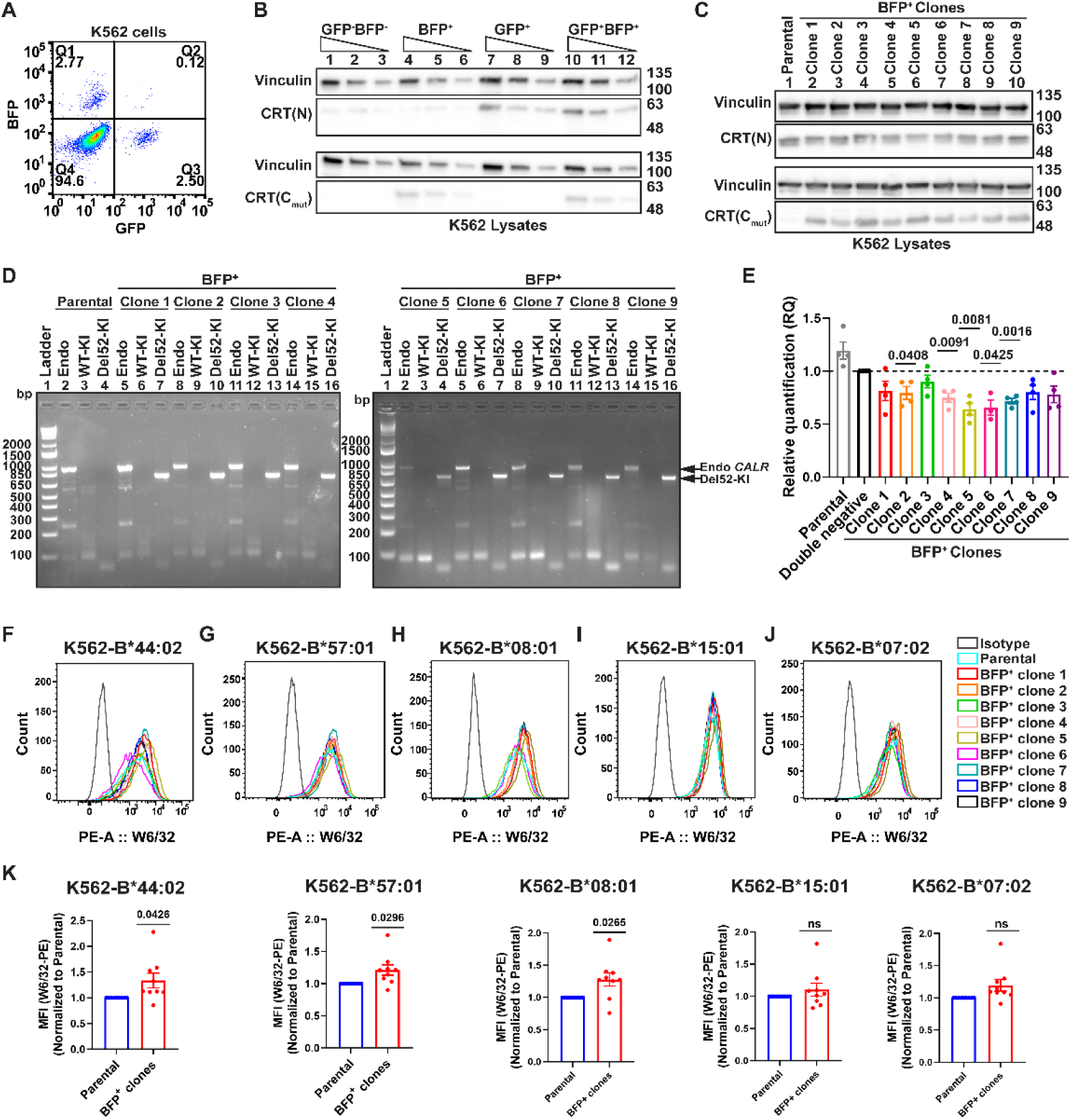
Calreticulin haploinsufficiency induces or maintains surface HLA class I expression for multiple allotypes. The CRISPR knock-in approach described earlier^34^ was used to insert wild-type or mutant *CALR* exon8 and exon9 sequences at the endogenous *CALR* locus. K562 cells were treated with Cas9/gRNA ribonucleoprotein (RNP) and recombinant adeno-associated virus serotype 6 (AAV6) for CRISPR/Cas9-mediated knock-in. Cells with successful knock-in of wild-type *CALR* and/or mutant *CALR* fragments were selected based on the expression of green fluorescent protein (GFP) and/or blue fluorescent protein (BFP), respectively. The reporter genes were inserted downstream of the respective knock-in fragments and driven by an independent spleen focus-forming virus (SFFV) promoter. **(A)** The dot plot shows the percentages of BFP^+^ (Q1), GFP^+^BFP^+^ (Q2), GFP^+^ (Q3), and GFP^-^BFP^-^ (Q4) K562 cells determined by flow cytometry following treatment of cells for CRISPR knock-in. **(B and C)** Representative immunoblots show wild-type CRT and mutant CRT expression detected using anti-CRT(N) and anti-CRT(C_mut_) antibodies, respectively, in the lysates of sorted GFP^-^BFP^-^, BFP^+^, GFP^+^ and GFP^+^BFP^+^ K562 cells (B) and various clones of BFP^+^ single cells compared to K562 parental cells (C). Vinculin is shown as the loading control. **(D)** Seamless integration of *CRT_Del52_*-specific knock-in fragment into the endogenous *CALR* locus was verified by in-out PCR using genomic DNA of indicated BFP^+^ single cell clones and compared to the K562 parental cells. Endogenous *CALR* (Endo), Del52 knock-in (Del52-KI), and wild-type *CALR* knock-in (WT-KI) fragments were amplified using a common forward primer and specific reverse primers. The agarose gel image shows PCR products amplified from endogenous *CALR* (Endo *CALR*) locus (∼919 bp) and *CRT_Del52_* knock-in (Del52-KI) allele (∼746 bp) as indicated. **(E)** RT-qPCR was performed to compare wild-type *CALR* mRNA expression in K562 parental cells, BFP^+^ clones, and K562 double negative cells (n=4). The expression levels of *GAPDH* and β*-actin* (n=3) or *GAPDH* and *HPRT* (n=1) were measured as endogenous reference controls. The primers specific to a target region within the wild-type *CALR* exon9 were used. **(F-J)** K562 parental cells and BFP^+^ clones 1 to 9 were transduced to express HLA-B*44:02 (E), HLA-B*57:01 (F), HLA-B*08:01 (G), HLA-B*15:01 (H), and HLA-B*07:02 (I). Representative histograms show surface staining with W6/32-PE or Isotype control (IgG2a-PE) antibody on the K562 parental cells or BFP^+^ clones. **(K)** Graphs show average W6/32 MFI value calculated from MFI values of individual BFP^+^ clones normalized to the MFI values of K562 parental cells from independent experiments (n=3). Graphs were plotted using GraphPad Prism, and statistical significance in panels E and K was determined using one-sample t-tests.

Nine single-cell clones of K562 GFP^-^BFP^+^ cells and one clone of THP-1 cells were used for further assessment of HLA class I expression after confirming the expression of both wild-type and mutant CRT proteins by immunoblots (**Figure 1C and Supplementary Figure 1A**). Seamless integration of the Del52 knock-in fragment at the correct site within the *CALR* gene was confirmed by PCR using genomic DNA isolated from the individual clones as a template. The BFP^+^ clones showed amplification from both endogenous *CALR* (∼919 bp) and *CRT_Del52_*knock-in locus (∼746 bp), unlike the endogenous *CALR* fragment alone observed for K562 parental (**Figure 1D**) or THP-1 GFP^-^BFP^-^ cells (**Supplementary Figure 1B**). Wild-type *CALR* mRNA expression was reduced in K562 BFP^+^ clones in comparison to the K562 parental or GFP^-^BFP^-^ cells, indicating CRT haploinsufficiency in cells with heterozygous Del52 knock-in (**Figure 1E**). Parental K562 cells and BFP^+^ clones were engineered to express five HLA-B allotypes individually; HLA-B*44:02, HLA-B*57:01, HLA-B*08:01, HLA-B*15:01, and HLA-B*07:02. HLA-B allotypes were chosen as this locus is the most polymorphic, and HLA-B allotypes show high variation in the requirements for other assembly factors for surface expression^25–27^. Surface HLA class I expression was measured by flow cytometry using a pan HLA class I-specific W6/32 antibody that binds to HLA class I-peptide complexes^35,36^. We observed small effects on the surface expression of all the HLA class I allotypes in BFP^+^ clones when compared to the K562 parental cells (**Figures 1F-K**) and THP-1 GFP^-^BFP^-^ cells **(Supplementary Figures 1C and D**). Compared to K562 parental cells, the K562 BFP^+^ clones exhibited a small but significant increase in the surface expression of HLA-B*44:02, B*57:01, and B*08:01, while the effects were non-significant for B*07:02 and B*15:01 surface expression (**Figure 1K**). Surface HLA class I levels were also slightly but non-significantly higher in the THP-1 BFP^+^ clone compared to the THP-1 GFP^-^BFP^-^ cells (**Supplementary Figures 1C and D**). Thus, heterozygous knock-in of MPN-linked CRT mutation at the endogenous *CALR* locus induces or maintains surface HLA class I expression in an allele-specific manner.

### Calreticulin dependencies of HLA class I allotypes and failure of MPN mutants to reconstitute expression in cells with full wild-type calreticulin deficiency

We could not isolate BFP^+^ clones with a homozygous knock-in of MPN-linked *CALR* mutation and thus adopted a different approach to measure the effects of full wild-type CRT deficiency on surface HLA class I expression. K562 cells were modified for stable expression of an expanded set of HLA class I allotypes, followed by knockdown of CRT expression in triplicate using CRISPR/Cas9 (**Supplementary Figure 2**). We observed an almost complete loss of CRT expression in the various K562-derived cell lines (**Supplementary Figure 2**). There was indeed allotype-dependent downregulation (CRT-dependency) of surface HLA class I expression following the CRT knockout when compared to the vector control cells (**Figure 2A**). Among the tested allotypes, HLA-B*57:01 and HLA-B*44:02 exhibited a >4.5-fold reduction in surface expression in CRT-KO cells compared with the vector control (**Figure 2B**). Other allotypes showed a smaller but significant reduction of surface expression following CRT-knockdown, including HLA-B*53:01 (3.1-fold), B*08:01(2.7-fold), and HLA-A*02:01 (2-fold) (**Figure 2B**). The least CRT-dependent among the tested allotypes were HLA-B*35:01, HLA-B*40:01, HLA-B*07:02, HLA-B*18:01, HLA-B*15:01, and HLA-B*45:01, each of which exhibited a <2-fold reduction of surface expression in CRT-KO cells (**Figure 2B**). Reconstitution of wild-type CRT expression (+CRT_WT_) (**Supplementary Figure 3**) restored surface HLA class I levels of all tested allotypes (**Figure 2C and Supplementary Figure 4**) to an extent largely consistent with their measured CRT dependencies (**Figure 2D**). However, the reconstitution of mutant CRT (CRT_Del52_ and CRT_Ins5_) expression failed to rescue surface expression of any of the tested allotypes (**Figures 2E, 2F and Supplementary Figure 4**). Thus, in contrast to CRT haploinsufficiency, complete CRT deficiency results in allotype-dependent reduction surface HLA class I expression. Furthermore, MPN mutants are unable to restore HLA class I expression in cells with full deficiency of wild-type CRT.

**Figure 2:**
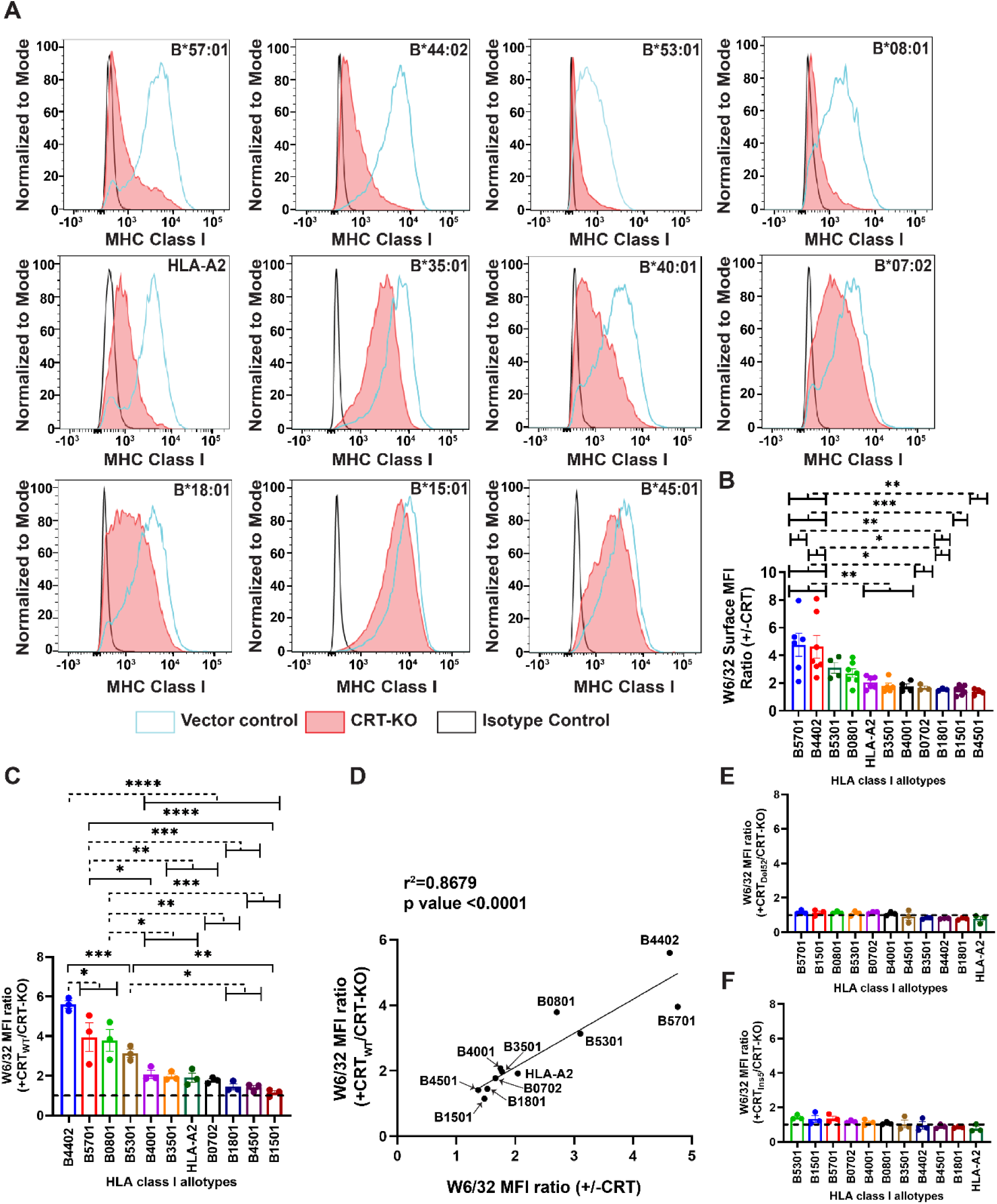
K562 cells with monoallelic HLA class I expression illustrate the allele-specific effects of calreticulin knockout on surface HLA class I expression and the failure of MPN mutants to reconstitute expression. HLA class I null K562 cells were transduced with retroviruses to express individual HLA class I allotypes and subsequently transduced with a pLentiCRISPRv2-based control virus (vector control) or those carrying CRT-targeting gRNA for CRISPR/Cas9-based knockdown of CRT expression (CRT-KO). K562 CRT-KO cells expressing individual HLA class I allotypes were further transduced for reconstitution of either wild-type CRT (CRT_WT_) or mutant CRT (CRT_Del52_ or +CRT_Ins5_) expression. Surface HLA class I expression was measured by flow cytometry following staining with the W6/32 antibody. **(A)** Representative histograms show surface staining of K562 cells expressing the indicated HLA class I allotypes with the W6/32 or the isotype control antibody. HLA class I expression is compared in the CRT-KO and vector control cells. **(B)** CRT-dependence of individual HLA class I allotypes is depicted as a ratio of surface HLA class I expression in vector control (+CRT) cells relative to that in CRT-KO cells (-CRT) quantified from flow cytometry experiments. Each dot shows the average MFI calculated from three biological replicates of CRT knockdown cells stained simultaneously in parallel experiments (n=3 for B*07:02 and B*18:01; n=4 for B*40:01, B*45:01 and B*53:01; n=5 for B*35:01; n=6 for B*57:01 and HA-A*02:01; n=7 for B*44:02, B*08:01 and B*15:01). **(C, E and F)** Graphs show MFI values of surface staining with the W6/32 antibody of individual HLA class I-expressing K562 CRT-KO cells reconstituted with either wild-type CRT (+CRT_WT_) (C) or Del52 (+CRT_Del52_) (E) or Ins5 (+CRT_Ins5_) (F) proteins. Data are plotted as a ratio of W6/32 MFI values of the cells reconstituted with CRT proteins relative to the MFI values from cells transduced with empty vector (CRT-KO). Each dot represents an individual biological replicate of reconstitution and all three biological replicates were stained in parallel experiments (n=3 for K562 cells-expressing B*07:02, B*18:01, B*40:01, B*45:01, B*53:01, B*57:01, and HA-A*02:01; n=5 for K562 cells-expressing B*08:01, B*15:01, B*35:01 and B*44:02). **(D)** Pearson correlation analysis between the loss of surface expression of indicated HLA class I allotypes upon CRT knockdown and the rescue of their surface expression following reconstitution of wild-type CRT expression. The loss or rescue of surface HLA class I expression is calculated as the ratio of the MFI value of surface staining with W6/32 antibody in vector control cells or CRT-KO cells reconstituted with CRT_WT_ relative to the MFI value of CRT-KO cells. The square of the Pearson correlation coefficient (r^2^) and p value are indicated on the graphs. Statistical significance in panels B, C, E and F was calculated in GraphPad Prism using ordinary one-way ANOVA analysis. * p value < 0.05, ** p value < 0.01,*** p value < 0.001. HLA, Human Leukocyte Antigen; MFI, Mean Fluorescent Intensity; MPN, Myeloproliferative Neoplasm

### Correlations of HLA class I CRT dependencies with the effects of CRT haploinsufficiency and with tapasin dependencies

Interestingly, HLA class I allotypes that exhibited a strong dependency on wild-type CRT (Figures 2B and 2C) also showed a significant induction of expression following heterozygous knock-in of Del52 mutation (**Figure 3A**). CRT, along with tapasin, is a component of the HLA class I assembly complex (**Figure 3B**). The CRT dependencies observed for individual HLA class I allotypes in K562 cells are positively correlated with the previously reported tapasin dependencies^25,27^ of these allotypes (**Figure 3C**). Although CRT and tapasin may have distinct roles within the PLC, HLA class I molecules appear to have different degrees of intrinsic (PLC-independent) *vs.* PLC-dependent assembly.

**Figure 3:**
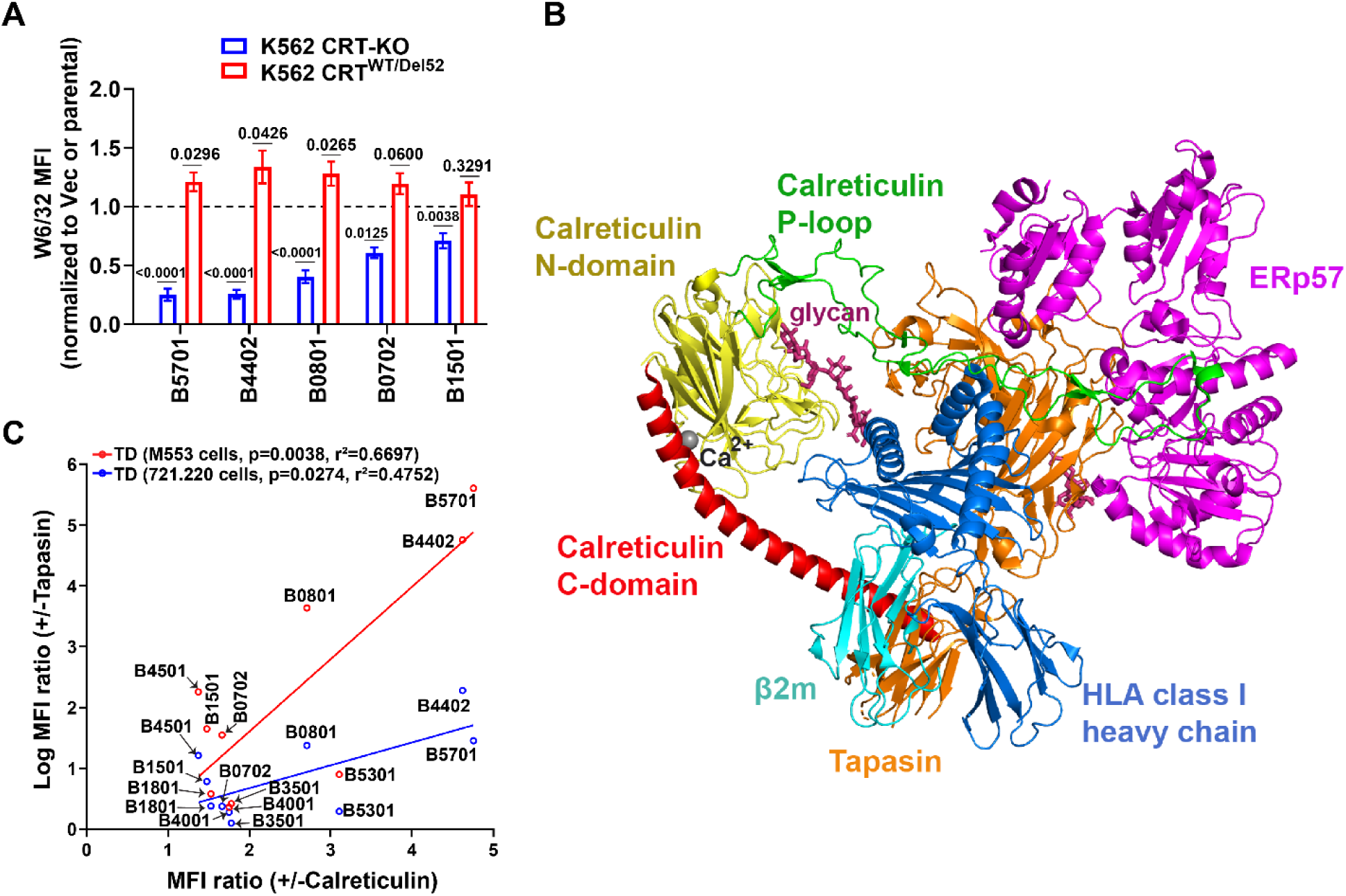
Correlations of HLA class I CRT dependencies with the effects of CRT haploinsufficiency and with tapasin dependencies. **(A)** The graph compares MFI ratios of surface staining with W6/32 antibody in cells with heterozygous CRT_Del52_ knock-in (CRT^WT/Del52^) (Figure 1K) and those with full wild-type CRT deficiency (CRT-KO) (Figure 2B). The ratios were calculated for CRT^WT/Del52^ cells or CRT-KO cells expressing the indicated HLA class I allotypes relative to K562 parental cells or vector control cells, respectively. Statistical significance was calculated using one-sample t-test in GraphPad Prism. **(B)** A model of HLA class I peptide loading complex (PLC) based on the cryo-EM structure (PDB ID: 7QPD)^49^ is shown. The individual protein components of PLC are shown in different colors (β2m in cyan; HLA class I heavy chain in blue; ERp57 in magenta; tapasin in orange). The globular domain of CRT, comprising 18-203 residues of the N-domain and 301-335 residues of the C-domain, is shown in yellow. The P domain of CRT, comprising residues 206-300, is shown in green color, and the acidic C-terminal helix, spanning residues 336-417, is shown in red color. The glycan moieties attached to the N86 residue of HLA class I heavy chain and the N233 residue of tapasin are shown in deep pink color and a calcium ion bound to the high-affinity calcium binding site D328 in the globular domain of CRT is shown as a grey sphere. **(C)** Pearson analyses were used to determine the correlation between CRT dependencies of the indicated HLA class I allotypes and their tapasin dependencies (TD) reported in M553 cells^25^ and 721.220 cells^27^. The dependencies of individual HLA class I allotypes on CRT and tapasin are calculated as the ratio of the MFI values of their surface expression in cells expressing CRT or tapasin, compared to the MFI values in cells lacking these respective chaperone proteins. The square of the Pearson correlation coefficient (r^2^) and p values are indicated on the graphs.

### Complete loss of wild-type CRT has small effects on the overall HLA class I expression in cell lines, and variable effects on the expression of individual allotypes

Most MPN patients have heterozygous loss of wild-type *CALR*, but some patients progress to homozygous loss of both wild-type *CALR* alleles^1,31^. Endogenous HLA class I expression in cells is achieved by co-dominant expression of at least six HLA class I alleles: two HLA-A’s, two HLA-B’s, and two HLA-C’s. Our findings thus far predict no losses in surface HLA class I expression in MPN patients with heterozygous expression of wild-type and mutant *CALR* alleles. Allele-dependent losses of HLA class I expression are predicted upon knockout of wild-type *CALR* and reconstitution with mutant *CALR*. However, the net effects on total HLA class I expression are expected to be averaged over the effects on individual HLA class I alleles expressed in the cells. The expression levels of HLA-C are known to be lower than HLA-A/B, and thus, HLA-C alleles would contribute less to the weighted average.

To further examine these predictions, we assessed the effects of CRT-KO and mutant CRT re-expression in a monocytic cell line and a B lymphoblastoid cell line (B-LCL) simultaneously expressing multiple HLA class I molecules. We used the pan HLA class I antibody, W6/32^35,36^ to assess the effects on total HLA class I, as well as the allele-specific antibodies to measure the expression levels of individual allotypes or combinations of allotypes. HLA-A*02:01 (HLA-A2) is a relatively frequent HLA class I allele that is detected using the BB7.2 antibody^37^. The specificity of this antibody for HLA-A2 was first verified using FlowPRA™ single antigen beads (One lambda). The binding analyses suggest that BB7.2 is largely specific for HLA-A*02:01 (**Figure 4A**), although weak binding to HLA-A*29:02, HLA-A*68:01, HLA-A*34:01, and HLA-A*31:01 was measured (**Figure 4B**). We also used the One Lambda monoclonal anti-Bw4 and anti-Bw6 antibodies for HLA-B/C protein measurements, based on previously established antibody specificities^38^.

**Figure 4:**
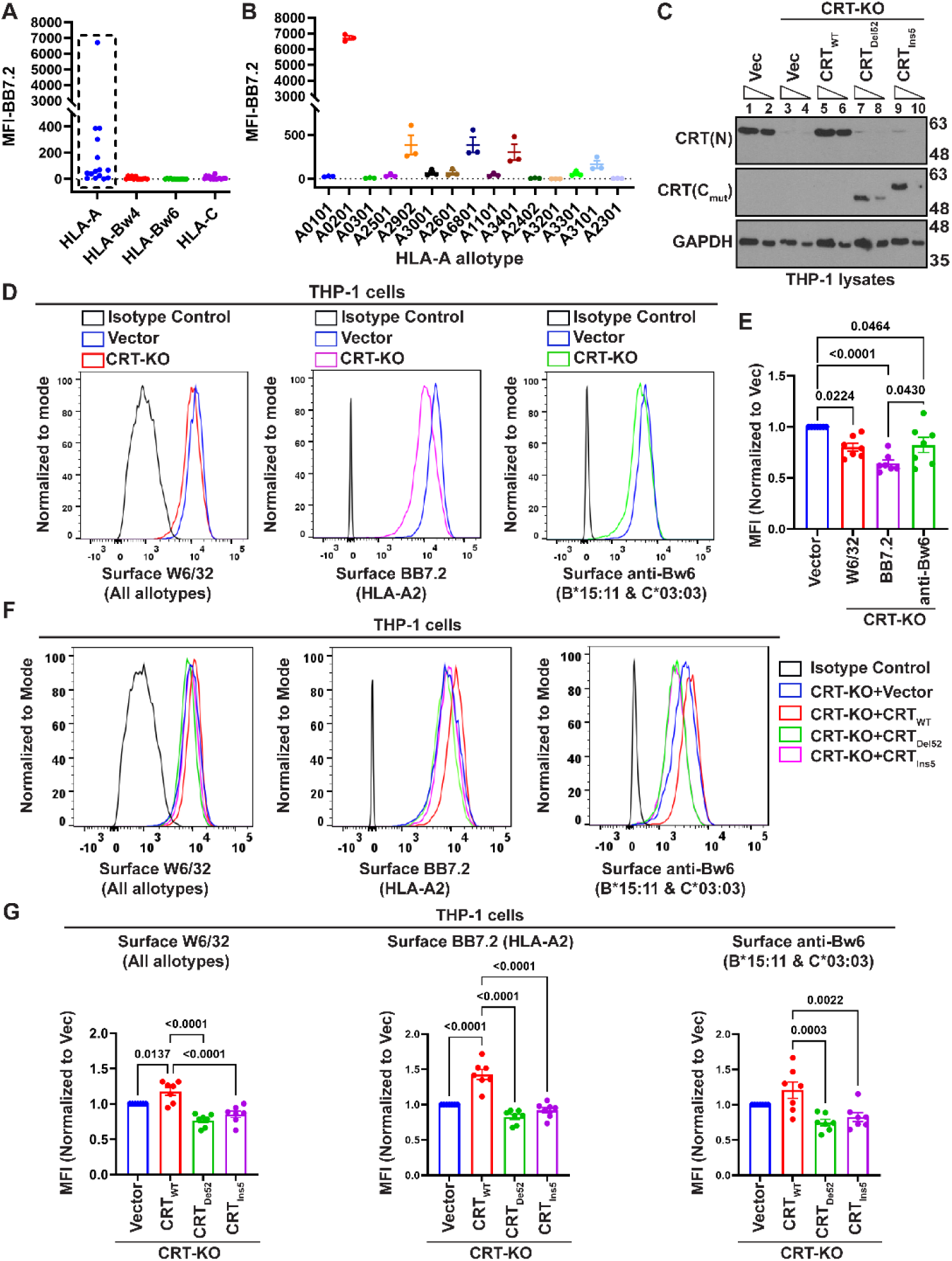
Allele-specific CRT dependencies of HLA class I expression in THP-1 cells, and failure of MPN mutants to support HLA class I expression. **(A and B)** FlowPRA™ single antigen HLA class I antibody detection tests (One lambda) were used to test the binding specificities of BB7.2 antibody to groups of beads coated with purified HLA class I single or fractionated antigens as indicated (A). The BB7.2 antibody binding to beads coated with individual HLA-A antigens is shown in panel B. Besides strong binding specificity for HLA-A*02:01, the BB7.2 antibody was also observed to show weaker non-specific binding to HLA-A*29:02, HLA-A*31:01, HLA-A*34:01 and HLA-A*68:01. **(C)** Representative immunoblots of the lysates of THP-1 cells (HLA-A*02:01/HLA*B*15:11/HLA-C*03:03; the published genotype for ATCC-derived cells^60^ was also verified as described in methods) subjected to CRISPR/Cas9-based knockdown of CRT expression and reconstitution of either wild-type CRT (CRT_WT_) or MPN-linked CRT mutants (CRT_Del52_ or CRT_Ins5_). Vec in lanes 1 and 2 represents THP-1 cells transduced with empty pLentiCRISPRv2 vector. Vec in lanes 3 and 4 refer to THP-1 CRT-KO cells transduced with empty vector during reconstitution. Two different amounts of each lysate were loaded in consecutive lanes. **(D and E)** Surface HLA class I levels in THP-1 cells were measured by flow cytometry using the W6/32 antibody (which detects HLA-A*02:01, HLA*B*15:11 and HLA-C*03:03 in THP-1 cells), anti-Bw6 (which detects B*15:11 and C*03:03 in THP-1 cells), and the HLA-A*02:01-specific BB7.2 antibody. Representative histograms (D) and averaged (n=7) MFI values of surface staining in CRT-KO cells normalized to the MFI values for THP-1 cells transduced with empty vector (E) are shown. **(F and G)** THP-1 CRT-KO cells reconstituted with either wild-type CRT (CRT-KO+CRT_WT_) or MPN-linked CRT mutants (CRT-KO+CRT_Del52_ or CRT-KO+CRT_Ins5_) or those transduced with empty vector (CRT-KO+Vector) were analyzed for surface HLA class I expression by flow cytometry using the W6/32, anti-Bw6, and BB7.2 antibodies. Representative histograms (F) and bar graphs with averaged MFI values (G) are shown. MFI data is normalized relative to the CRT-KO+Vector cells (n=7). Statistical significance was calculated using one-way ANOVA for panels E and G. Graphs are plotted in GraphPad Prism.

Surface HLA class I levels were measured in THP-1 cells (ATCC; HLA-A*02:01/HLA-B*15:11/HLA-C*03:03) following CRISPR/Cas9-based CRT knockdown (**Figure 4C**). There was a 1.25-fold reduction in HLA-B*15:11/HLA-C*03:03 expression, detected by anti-Bw6 antibody in THP-1 CRT-KO cells compared to vector control cells. Surface HLA-A*02:01 expression was reduced by 1.6-fold in THP-1 CRT-KO cells compared to vector control cells (**Figures 4D and 4E**). In combination, the overall surface HLA class I expression change measured with the W6/32 antibody was about 1.25-fold, more similar to that measured with anti-Bw6 than with the BB7.2 antibody, likely due to the contributions of two HLA class I allotypes to the anti-Bw6 signal. As expected, the reconstitution of wild-type CRT expression (CRT_WT_) (**Figure 4C**) restored surface HLA class I levels, but the MPN-linked CRT mutants (CRT_Del52_ or CRT_Ins5_) failed to rescue surface HLA class I expression in THP-1 CRT-KO cells for all three allotypes (**Figures 4F and 4G**).

CRT expression was also targeted by CRISPR/Cas9 (**Supplementary Figures 5A and 5B**) in the IHW01147 B-LCL expressing HLA-A*01:01/A*02:01, HLA-B*08:01/B*44:02 and HLA-C*07:01/C*05:01. Surface HLA-A*02:01 levels were reduced by 1.5-fold in IHW01147 CRT-KO cells relative to the vector control cells. Surface staining with anti-Bw6, which detects the expression of HLA-B*08:01 and C*07:01 on the IHW01147 B-LCL, also indicated a 1.5-fold reduction. Consistent with the K562 data, surface HLA-B*44:02 expression, detected with the anti-Bw4 antibody, indicated a more significant downregulation (∼4-fold) in CRT-KO cells compared to vector control cells (**Supplementary Figures 6A and 6B**). The overall class I surface expression, measured using the W6/32 antibody, was reduced by about 1.3-fold following the CRT knockdown, which was higher than predicted by measurements with allele-specific antibodies, but nonetheless lower than in cells that retain CRT expression (**Supplementary Figures 6A and 6B**). It is possible that the conformational preferences of the allele-specific antibodies overestimate the effects of CRT knockdown on surface HLA class I expression. Reconstituting wild-type CRT expression in IHW01147 CRT-KO cells (**Supplementary Figure 5C**) restored surface HLA class I levels (**Supplementary Figures 6C and 6D**). We observed a robust increase in surface HLA-B*44:02 expression (7.2-fold) in IHW01147 CRT-KO cells upon reconstitution of wild-type CRT expression (**Supplementary Figure 6D**). Again, as expected, mutant CRT proteins (CRT_Del52_ and CRT_Ins5_) in IHW01147 CRT-KO cells (**Supplementary Figure 5C**) failed to restore surface HLA class I levels (**Supplementary Figures 6C and 6D**). Overall, the findings from human cell lines expressing multiple HLA class I allotypes confirmed allele-specific effects of CRT-KO upon surface HLA class I expression and small but significant negative effects on combined HLA class I expression.

### Surface HLA class I expression in platelets from MPN patients with *CALR* mutations is generally in the normal range

Based on the studies thus far, we predicted small effects of *CALR* mutations upon the overall surface HLA class I expression in MPN patient-derived cells, to an extent dependent on their HLA allelic make-up and homozygous *vs*. heterozygous *CALR* mutations. As noted above, *CALR* mutations generally present heterozygously in MPN, with one copy each of wild type and mutant *CALR* and a small number of *CALR*-MPN patients progress to homozygosity^1,31^. To identify patients expressing homozygous *CALR* mutations in blood cells, we performed immunoblots of platelet lysates, where available (**Figure 5A**). Alternatively, or additionally, we estimated variant allele frequency (VAF) using genomic DNA purified from patient peripheral blood mononuclear cells (PBMC). Clinical VAF values were also used where VAF measurements on PBMC were not available (**Supplementary Table 1)**.

**Figure 5:**
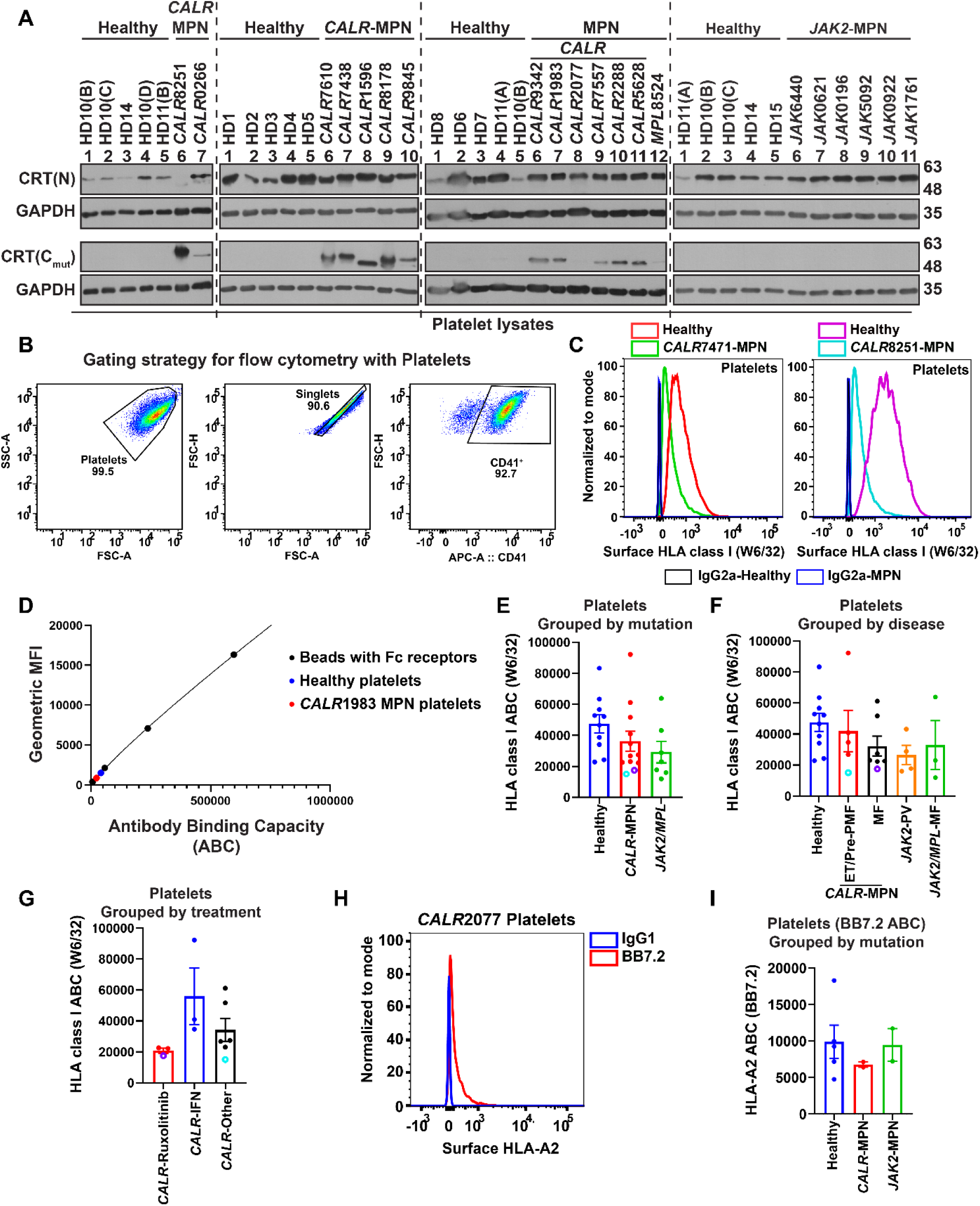
Surface HLA class I expression in platelets from MPN patients with *CALR* mutations generally falls within the normal range. **(A)** Immunoblots show wild-type and mutant CRT expression in the lysates of platelets purified from blood samples of healthy donors (HD) and MPN patients with *CALR*, *JAK2,* or *MPL* mutations as indicated. Each lane represents an individual patient or healthy donor sample. The wild-type and mutant CRT proteins were detected using anti-CRT(N) and anti-CRT(C_mut_) antibodies, respectively. GAPDH was detected as a loading control. Quantitative flow cytometry was used to measure surface HLA class I expression on platelets purified from MPN patients or healthy donors after staining with W6/32 or BB7.2 antibodies or their respective isotype controls. **(B)** The dot plots show the gating strategy used for flow cytometry-based identification of CD41^+^ platelets. **(C)** Representative histograms show surface staining with the W6/32 antibody or the IgG2a isotype control, as detected by flow cytometry, on CD41^+^ platelets isolated from *CALR*-MPN patients, *CALR*7471 and *CALR*8251, compared to the healthy donor platelets prepared on the same day. **(D)** Beads with Fc receptors (Quantum simply cellular anti-mouse beads, Bangs Laboratories, Inc.) were stained simultaneously with the platelets using the same antibody dilution to determine the Antibody Binding Capacity (ABC) of platelets. Standard curves were generated by plotting numbers of Fc receptors (provided by the manufacturer) against the geometric mean fluorescence intensity (gMFI) values measured for individual bead populations. The representative standard curve shows data points corresponding to the gMFI values of healthy donor (blue) and MPN patient (*CALR*1983, red) platelets along with those for the individual bead populations (black). (**E-G**) Graphs show W6/32 antibody binding capacities of platelets from healthy donors (n=10) and MPN patients. The MPN patient samples were grouped as those with *CALR* mutations (*CALR*-MPN; n=12) or *JAK2/MPL* mutations (n=6 for *JAK2* and n=1 *MPL* mutation) (E). The *CALR*-MPN patients were further grouped based on the disease into ET/Pre-PMF (n=5) and MF (n=7) groups, and the patients with *JAK2/MPL* mutations were grouped into PV (n=4) and MF (n=3) groups (F). The *CALR*-MPN patients were grouped into *CALR*-Ruxolitinib (n=3), *CALR*-IFN (n=3) and *CALR*-other (n=6) groups based on their clinical treatments (G). Each dot represents an individual blood donor, and average ABC values are plotted for multiple collections from the same healthy or patient donor. MPN patients with evidence for homozygous *CALR* mutations (Supplementary Table 1), *CALR*8251 (purple) and *CALR*7471 (cyan), are highlighted in the graphs. **(H)** Representative histogram shows surface staining with BB7.2 antibody on platelets from HLA-A2^+^ MPN donor *CALR*2077 compared to the IgG2a isotype control antibody staining. **(I)** The graph shows the platelet BB7.2 ABC values from healthy donors (n=5), MPN patients with *CALR* mutations (*CALR*-MPN; n=2), and MPN patients with *JAK2* mutations (*JAK2*-MPN; n=2). All graphs were plotted in GraphPad Prism and show the mean values, while the error bars show the standard error of the mean. Statistical significance was calculated using ordinary one-way ANOVA for panels E to G and I. ET, Essential Thrombocythemia; MF, Myelofibrosis; PMF, Primary Myelofibrosis; IFN, Interferon

For these analyses, platelets were purified from fresh blood samples of MPN patients with *CALR* or *JAK2* mutations and healthy controls. Mutant CRT expression was detected by immunoblots with the anti-CRT(C_mut_) antibody^8,39^ in platelet lysates from *CALR*-MPN patients, but not in healthy donors (HD) or JAK2 patients (**Figure 5A**). A faint band was also detected with the anti-CRT(C_mut_) antibody in platelet lysates from one patient with *MPL*W515K mutation, likely due to a non-specific signal. We also detected wild-type CRT expression in platelet lysates from all MPN patients except *CALR*8251, which showed a strong band with anti-CRT(C_mut_) antibody and undetectable wild-type CRT levels (**Figure 5A**). The *CALR* VAF in PBMC for this patient was estimated at 91.3% (**Supplementary Table 1**). Taken together with the immunoblotting data, this VAF is consistent with the presence of cells with homozygous *CALR* mutations in this patient. Another patient, *CALR*7471, had a clinically-derived whole blood VAF of 76.2%, which indicated the presence of cells with homozygous *CALR* mutations. Four additional patients had *CALR* VAF>20 in PBMC (**Supplementary Table 1**). Among PBMC subsets, monocytes have the highest mutant *CALR* VAF^40^, but comprise only 5-10% of PBMC in normal individuals. A PBMC VAF>20 could thus be indicative of the presence of hematopoietic cells with homozygous mutations; however, both wild-type and mutant CRT proteins were detected in platelet lysates of these four patients (**Figure 5A**), consistent with the prevalence of cells with heterozygous mutations. Thus, among the 16 *CALR*-MPN patients whose HLA class I expression levels were measured, it appears that only two patients, *CALR*8251 and *CALR*7471, have progressed to homozygosity.

Surface HLA class I levels were measured by quantitative flow cytometry using the pan-HLA class I W6/32 antibody. This was undertaken in platelets and PBMC, with both samples included for most patients. Platelets were gated for high CD41 expression (**Figure 5B**) as shown earlier^39^. Representative histograms show W6/32 staining of platelets from MPN patients with *CALR* mutations relative to the healthy donor platelets processed simultaneously with the patient samples (**Figure 5C**). W6/32 binding to beads with known Fc receptor levels was measured in parallel with the platelet analyses. This enabled the conversion of cell-derived W6/32 geometric mean fluorescence intensities (gMFI) to antibody binding capacities (ABC), allowing quantitative comparisons of HLA class I levels across samples collected over several months (**Figure 5D**). As previously described for leukocytes^38^, there is a considerable donor-to-donor variation in HLA class I expression even among the healthy donor platelets, ranging from ABC values of 22905 to 83400. The platelet W6/32 ABC values for MPN patients with *CALR* mutations fall within the range of healthy donor ABC values as well as those derived for platelets from MPN patients with *JAK2*/*MPL* mutations (**Figure 5E and Supplementary Table 1**). There were also no significant differences when *CALR*-MPN patients were grouped by disease or treatment regimens. Notably, surface HLA class I levels on platelets from both *CALR*8251 and *CALR*7471 were at the lower end of the measured range (**Figures 5E-G**).

With the relatively small cohort size, high degree of HLA polymorphisms, and cross-reactivity patterns of anti-HLA-Bw4 and anti-HLA-Bw6, that we have previously used for allele-specific HLA class I measurements^38^, allele-specific expression comparisons were only possible for HLA-A2. The BB7.2 antibody was used to compare HLA-A2 expression on platelets (**Figure 5H**) from HLA-A2^+^ MPN patients and healthy donors (**Supplementary Table 2**; excluding samples that co-expressed another BB7.2-reactive HLA-A (Figure 4B) or any HLA-A allotype not included within the FlowPRA screening). HLA-A2 expression on platelets from MPN patients with *CALR* mutations was within the range found in healthy donors and MPN patients with *JAK2* mutations (**Figure 5I**). Overall, total HLA class I levels on platelets in most *CALR*-MPN patients fall within normal ranges; however, patients with evidence for the presence of homozygous *CALR* mutations have expression levels that fall at the lower end of the expression range.

### Classical monocytes display HLA expression levels responsive to treatment type

Among PBMC, monocytes have the highest VAF for *CALR* mutations^40^, and classical monocytes (CD3^-^CD19^-^HLA-DR^+^CD14^+^CD16^-^) are generally the most abundant among monocyte subsets (**Figure 6A**). Mutant CRT proteins lack the ER-retention KDEL motif and are trafficked to the cell surface^5,10,39,41,42^. The surface expression of mutant CRT has been previously measured on monocytes from MPN patients^41^. By flow cytometry, we generally detected higher surface staining with both anti-CRT(N) and anti-CRT(C_mut_) antibodies in classical monocytes from MPN patients compared to healthy donor samples collected on the same day (representative histograms for monocytes derived from *CALR*2077 are shown in **Figure 6B**). Although most patients showed higher anti-CRT surface staining on classical monocytes compared with same-day healthy donors, higher surface CRT was measured on the classical monocytes from MF patients compared to ET/Pre-PMF patients (**Figure 6C**), suggesting that MF may induce changes to CRT expression levels. Notably, *CALR*0387 and *CALR*2077, who are PMF patients, had high surface CRT expression on their classical monocytes despite low VAF in PBMC (Supplementary Table 1). Low monocyte fractions were detected among live cells in PBMC by flow cytometry for both these patient samples, which could contribute to their low VAF, as monocytes have the highest *CALR* VAF among PBMC subsets^40^.

**Figure 6:**
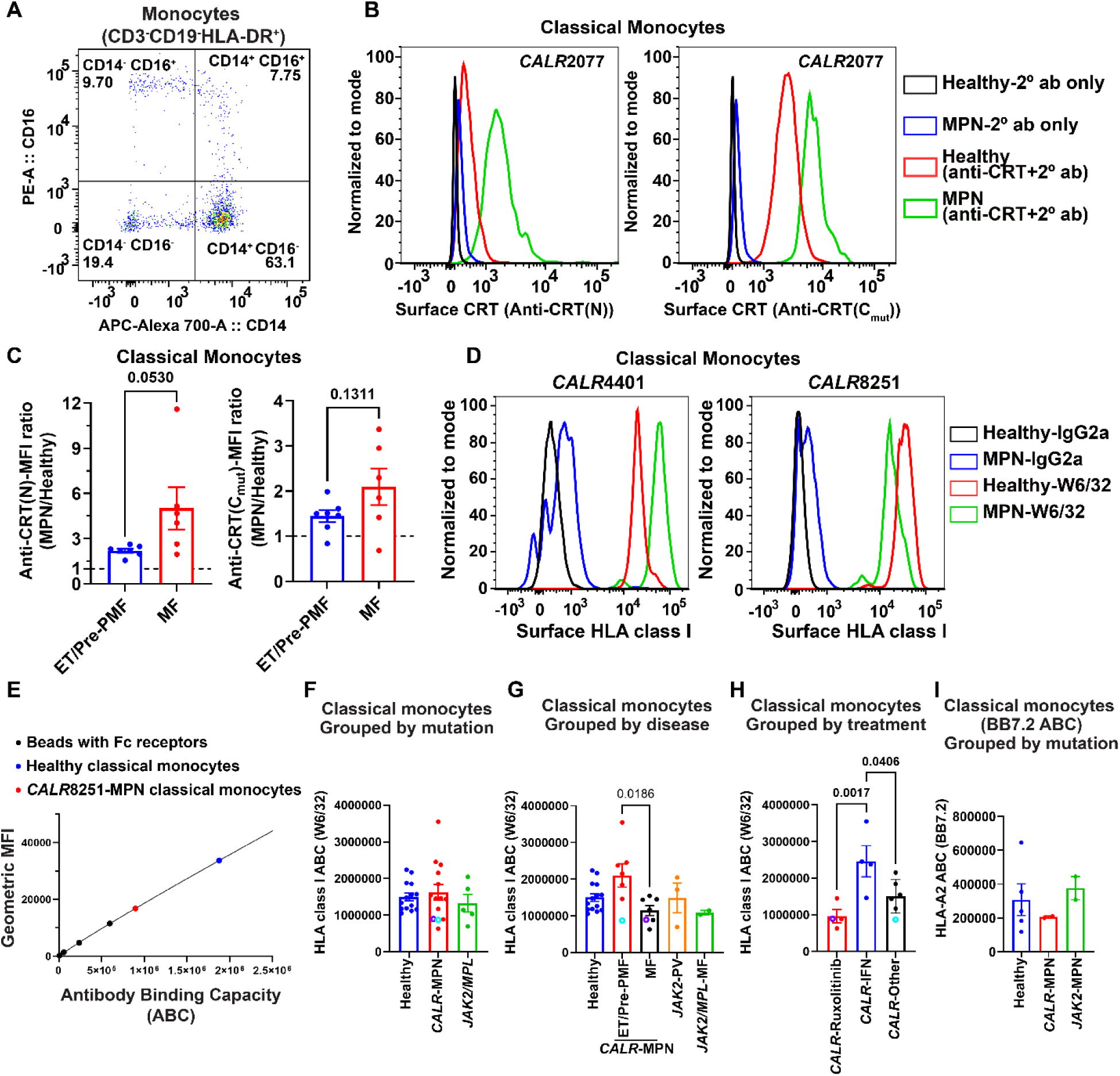
Mutant CRT and HLA class I expression in classical monocytes from healthy donors and MPN patients. Peripheral blood mononuclear cells (PBMC) isolated from blood samples of MPN patients or healthy donors were stained for monocyte-specific surface markers to assess surface CRT or surface HLA class I expression. **(A)** Representative dot plot shows gating of CD3^-^CD19^-^HLA-DR^+^ monocytes based on the expression of CD14 and CD16 markers into classical (CD14^+^CD16^-^), intermediate (CD14^+^CD16^+^) and non-classical (CD14^-^CD16^+^) monocytes. **(B)** Representative histograms show surface CRT expression on classical monocytes in PBMC isolated from a *CALR*-MPN patient, *CALR*2077, and the same-day healthy donor, measured by flow cytometry using anti-CRT(N) or anti-CRT(C_mut_) primary antibody and Alexa Fluor 488-cojugated secondary antibody (2° ab). Surface staining with secondary antibody alone is shown as a control for both the healthy donor and the MPN patient. **(C)** The graphs show the mean fluorescence intensity (MFI) values of staining with anti-CRT(N) or anti-CRT(C_mut_) antibody on classical monocytes from MPN patients grouped as ET/Pre-PMF (n=7) and MF (n=6). These values are presented as a ratio relative to the MFI values of surface CRT staining of healthy donor cells that were processed on the same day as patient samples. **(D)** Representative histograms show surface staining with the W6/32 antibody or the IgG2a isotype control, as detected by flow cytometry, on classical monocytes in PBMC prepared from *CALR*-MPN patients, *CALR*4401 and *CALR*8251, compared to the staining of classical monocytes in healthy donor PBMC prepared on the same day as patient samples. **(E)** Beads with Fc receptors (Quantum simply cellular anti-mouse beads, Bangs Laboratories, Inc.) were stained simultaneously as the PBMC using the same antibody dilution to determine the Antibody Binding Capacity (ABC) of cells. Standard curves were generated by plotting numbers of Fc receptors (provided by the manufacturer) against the geometric mean fluorescence intensity (gMFI) values measured for individual bead populations. The representative standard curve shows data points corresponding to the gMFI values of classical monocytes from a healthy donor (blue) and MPN patient *CALR*8251 (red), as well as data points for individual bead populations (black). (**F-H**) Graphs show W6/32 ABC values of classical monocytes from healthy donors (n=14) and MPN patients. The MPN patient samples were grouped as those with *CALR* mutations (*CALR*-MPN; n=14) or *JAK2/MPL* mutations (n=4 for *JAK2* mutation and n=1 for *MPL* mutation) (F). The *CALR*-MPN patients were further grouped based on the disease as ET/Pre-PMF (n=7) and MF (n=7), and the patients with *JAK2/MPL* mutations were grouped into PV (n=3) and MF (n=2) groups (G). The *CALR*-MPN patients were also grouped into *CALR*-Ruxolitinib (n=4), *CALR*-IFN (n=4) and *CALR*-other (n=6) groups based on their clinical treatments (H). Each dot represents an individual blood donor, and average ABC values are plotted for multiple collections from the same healthy or patient donors. MPN patients with evidence for homozygous *CALR* mutations (Supplementary Table 1), *CALR*8251 (purple) and *CALR*7471 (cyan) have been highlighted in the graphs. **(I)** The graph shows the BB7.2 ABC values of classical monocytes from healthy donors (n=5), MPN patients with *CALR* mutations (*CALR*-MPN; n=2) and MPN patients with *JAK2* mutations (*JAK2*-MPN; n=2). All graphs were plotted in GraphPad Prism and show mean values, and the error bars show the standard error of the mean. Statistical significance was calculated using unpaired t-tests for panel C and ordinary one-way ANOVA for panels F-I. ET, Essential Thrombocythemia; MF, Myelofibrosis; PMF, Primary Myelofibrosis; IFN, Interferon

Using quantitative flow cytometry with the W6/32 antibody, we measured surface HLA class I levels on classical monocytes from MPN patients and healthy donors. Representative histograms show staining with W6/32 antibody on classical monocytes from *CALR*-MPN patients, *CALR*4401 and *CALR*8251, compared to classical monocytes from healthy donors processed simultaneously as patient samples (**Figure 6D**). The W6/32 ABC values for classical monocytes were determined by parallel staining of beads with known Fc receptor levels for the quantitative HLA class I measurements (**Figure 6E**). The W6/32 ABC values for classical monocytes from MPN patients with *CALR* mutations fall within a similar range as the ABC values for classical monocytes from healthy donors and MPN patients with *JAK2/MPL* mutations. Additionally, surface HLA class I levels on classical monocytes from *CALR*8251 and *CALR*7471 were at the lower end of the HLA class I expression range in monocytes, as was noted for platelets (**Figure 6F and Supplementary Table 1**). Moreover, ET/Pre-PMF samples exhibited significantly higher surface HLA class I expression in classical monocytes compared to MF samples (**Figure 6G**). Furthermore, significantly higher surface HLA class I levels were measured on classical monocytes from *CALR*-MPN patients treated with type-1 interferon compared to *CALR*-MPN patients in alternate treatment groups (**Figure 6H**). Type-1 interferon is a well-known inducer of MHC class I expression and of components of the MHC class I assembly pathway^43,44^, which could explain these observed trends. No significant differences were detected in HLA-A2 expression with BB7.2 antibody on classical monocytes from HLA-A2^+^ *CALR*-MPN patients when compared to healthy donors or *JAK2*-MPN patients (**Figure 6I**).

## Discussion

Downregulation of surface HLA class I expression is a strategy evolved by cancer cells to elude immune activation^45^. Here, we looked at the effects of *CALR* mutations upon surface HLA class I expression variations in cell lines and MPN patient platelets and monocytes. Most MPN patients have a high prevalence of hematopoietic clones with heterozygous *CALR* mutations^1,31^. We used a CRISPR-based approach^34^ to generate a heterozygous knock-in of the Del52 mutation at the endogenous *CALR* locus. Cells with heterozygous CRT_Del52_ expression did not show loss of surface HLA class I expression (**Figure 1**). These findings indicate that the wild-type CRT expression driven by a single non-mutated *CALR* allele is sufficient to maintain the HLA class I assembly and expression. In fact, the more CRT-dependent HLA class I allotypes show a small but significant increase in surface expression following the knock-in of one copy of *CRT_Del52_* allele, suggesting that CRT-dependent quality control normally restrains expression of such allotypes. These findings are consistent with prior findings of minimal or no effects on overall glycoprotein abundance and structural stability in granulocytes from patients with heterozygous *CALR* mutations^46^.

Homozygous *CALR* mutations are rare and are associated with type-2 mutations and disease progression^40,47^. As a component of the PLC, CRT interacts via its glycan binding site with the monoglucosylated glycan moiety^12,48,49^ attached to the highly conserved Asn 86 residue of the HLA class I heavy chain within the ER^17,50^. Sub-optimal peptide-MHC class I complexes with low stability were observed in CRT-deficient cells^11,30^. Complete loss of wild-type CRT expression was achieved using the standard CRISPR/Cas9-based gene knockout methodology (**Figures 2 and 4**), and this was followed by reconstitution with gRNA-resistant versions of different *CALR* constructs to simulate the homozygous *CALR* mutations in MPN patients. HLA-B*44:02, HLA-B*57:01, and HLA-B*53:01 showed the highest CRT dependencies (**Figure 2**). Similar trends of high dependence on other PLC components, tapasin and TAP, have been observed earlier for HLA-B*44:02 and HLA-B*57:01^25,27^. Within the PLC, in addition to the glycan-dependent CRT-MHC class I interactions, there are direct interactions between CRT and the thiol oxidoreductase ERp57. The tip of the arm-like P-domain of CRT binds ERp57^17,51^ which is, in turn, covalently linked to tapasin via a disulfide bond between the Cys95 of Tapasin and Cys57 of ERp57^16,52^. Tapasin also bridges β2m-bound MHC class I, CRT, and ERp57 to TAP^50^. This intricate but highly orchestrated network of interactions between the components of the PLC^14,16^ facilitates the assembly of HLA class I proteins with peptides and allows for optimization of the peptide loading. HLA class I molecules have varying degrees of dependency on PLC components for their assembly *vs.* intrinsic PLC-independent assembly, and the degree of dependency on CRT *vs.* tapasin is correlated (**Figure 3C**). HLA-B*53:01 is an exception since it exhibited a 3.1-fold loss of surface expression in CRT-deficient K562 cells in contrast to much smaller effects under tapasin-deficient conditions. The differences in the tapasin versus CRT dependence of HLA-B*53:01 indicate the existence of possible PLC-independent mechanisms of CRT-mediated stabilization of surface expression for some HLA class I allotypes.

While the re-expression of wild-type CRT restored the expression of HLA class I allotypes in CRT-KO cells, MPN-linked CRT_Del52_ and CRT_Ins5_ mutants failed to rescue surface HLA class I expression in CRT-deficient cells for any of the tested allotypes (**Figures 2 and 4**). In a previous study, the reconstitution of CRT_Del52_ or CRT_Ins5_ expression failed to restore surface MHC class I expression in CRT^null^ HEK293T cells^30^. The negatively charged C-terminal helix of CRT lies close to a positively charged region in the C-domain of tapasin^17^. The CRT_Del52_ and CRT_Ins5_ mutants have reduced association with tapasin in CRT^null^ cells, but the interactions were partially restored upon the addition of the KDEL motif to the C-terminus of the mutant proteins. However, the KDEL-mediated increased cellular retention of CRT_Del52_ and CRT_Ins5_ was insufficient for the recovery of surface MHC class I expression^30^. Although the mechanistic details of impaired chaperone activity remain to be fully elucidated, it is clear that the *CALR* mutations impede the role of PLC in catalyzing optimal peptide assembly onto HLA class I proteins.

The opposing effects of heterozygous vs. homozygous mutations of *CALR* on HLA class I expression (**Figure 3A**) might be explained by multiple roles of CRT in HLA class I assembly both within and outside of the PLC. Outside of the PLC, CRT functions in the retrieval of suboptimally assembled MHC class I molecules from ER-Golgi intermediate compartments to ER^53^. In the full absence of CRT, the loss of PLC function would cause impairments in both assembly and retrieval, leading to the expression of more unstable peptide-HLA class I complexes at the cell surface, and a net loss in the surface HLA class I expression. Under conditions of CRT haploinsufficiency, the PLC may be functional to mediate assembly of HLA class I proteins that are at least partly optimized, whereas the lower expression of retrieval-competent CRT might allow for more surface expression of these partly optimized HLA class I proteins. However, this effect is specifically observed for the CRT-dependent allotypes. CRT-independent allotypes are less impacted by both haploinsufficiency and the complete loss of wild-type CRT expression.

Consistent with cell line-based experiments, we did not observe major differences in surface HLA class I expression on platelets and classical monocytes from patients with *CALR* mutations when compared to healthy donors or patients with *JAK2/MPL* mutations (**Figures 5E and 6F**). Amongst ET/Pre-PMF samples, *CALR*7471 with a clinical VAF of 76.2%, showed among the lowest surface HLA class I expression on both platelets and classical monocytes (**Supplementary Table 1**). Low ABC values were also measured for *CALR*8251, a post-ET MF patient, who had a VAF of 91.3% in PBMC measured with a sample collected six years and 11 months before the surface HLA class I measurements. Both patients have at least one strongly CRT-dependent HLA-B (B*44:02 for *CALR*7471 and B*08:01 for *CALR*8251) (**Supplementary Table 2**). Since a sufficient number of MPN patients and healthy donors with matched HLA genotypes were unavailable, meaningful allele-level comparisons of HLA class I expression were not possible. Nonetheless, the low overall HLA class I expression confirms the negative effects of homozygous *CALR* mutations upon HLA class I expression in freshly isolated primary cells.

An analysis of *CALR*-MPN patients, categorized by disease, revealed a significant difference between ET and MF patients (**Figure 6G**). Further grouping of *CALR* patients by treatment indicated higher expression in patients treated with type-I interferon (**Figure 6H**). The type-I interferon treatment is prescribed as a cytoreductive therapy to *CALR*-ET patients and has been observed to mediate complete hematologic remission in patients^54,55^. Based on our results, we can hypothesize that the protective effects of type-I IFN treatment in MPN patients may, in part, be contributed to by the increased HLA class I-mediated antigen presentation (reviewed in^56^). A significant increase in the expression of HLA class I and TAP genes was measured in the whole blood samples from MPN patients after three months of interferon-alpha 2 treatment^57^. Another recent study reported no reduction of surface HLA class I and class II expression on PBMC and CD34^+^ cells from mutant *CALR*-positive MPN patients in comparison to the healthy donor cells. There was rather a significant increase in surface HLA class I expression on both PBMC and CD34^+^ cells from *CALR*-MPN patients when compared to healthy control cells (Figure S3 from^58^), which was also potentially attributed to interferon treatment of some patients.

It is also noteworthy that HLA class I ABC is low in MPN patients with MF, particularly those treated with Ruxolitinib (**Supplementary Table 1**). MF is associated with more severe disease phenotype and poorer prognosis among MPN patients with *CALR* mutations^59^. MF samples in our cohort include both primary myelofibrosis and post ET-MF samples. Overall, the impact of MPN-linked CRT mutations on surface HLA class I expression is complex, influenced by mutant allele burden, type of disease, and type of treatment. Nonetheless, the maintenance of surface HLA class I expression in the majority of MPN patients with heterozygous expression of CRT mutants offers hope for activation of protective immune responses against antigenic peptides derived from the mutated CRT sequence.

## Acknowledgments

We extend our gratitude to all the healthy and patient donors who volunteered to donate blood for this study. We also appreciate the support of Amanda Prieur and the University of Michigan Platelet Physiology and Pharmacology Core in collecting healthy donor blood samples. We also thank Polk Avery from Dr. Moshe Talpaz’s laboratory for coordinating the collection of blood samples from MPN patients. We acknowledge the University of Michigan vector core and flow cytometry core for assistance with CRISPR Knock-in experiments. We acknowledge Oloche Owoicho for providing THP-1 cells with calreticulin knockout and Fawziyah Khatri for helping with the generation of THP-1 CRT_Del52_ knock-in cells. This work was funded by the National Institute of Health grant (R01 AI123957) to MR and the University of Michigan Fast Forward Protein Folding Diseases Initiative.

## Author Contributions

AK and HD designed and performed experiments and analyzed the data. GP coordinated the collection of healthy donor blood samples, and MK coordinated the collection of blood samples from MPN patients. MT is the director of the MPN repository at the University of Michigan. AK wrote the original manuscript draft, and AK and GP edited the manuscript. MR designed and supervised the study, obtained funding, analyzed data, and wrote and edited the manuscript.

## References

1. Klampfl T, Gisslinger H, Harutyunyan AS, et al. Somatic Mutations of Calreticulin in Myeloproliferative Neoplasms. N Engl J Med. 2013;369(25):2379–2390.

2. Nangalia J, Massie CE, Baxter EJ, et al. Somatic CALR mutations in myeloproliferative neoplasms with nonmutated JAK2. N Engl J Med. 2013;369(25):2391–2405.

3. Baksh S, Michalak M. Expression of calreticulin in Escherichia coli and identification of its Ca2+ binding domains. J Biol Chem. 1991;266(32):21458–21465.

4. Wijeyesakere SJ, Gafni AA, Raghavan M. Calreticulin is a thermostable protein with distinct structural responses to different divalent cation environments. J Biol Chem. 2011;286(11):8771–8785.

5. Araki M, Yang Y, Masubuchi N, et al. Activation of the thrombopoietin receptor by mutant calreticulin in CALR-mutant myeloproliferative neoplasms. Blood. 2016;127(10):1307–1316.

6. Chachoua I, Pecquet C, El-Khoury M, et al. Thrombopoietin receptor activation by myeloproliferative neoplasm associated calreticulin mutants. Blood. 2016;127(10):1325–1335.

7. Elf S, Abdelfattah NS, Chen E, et al. Mutant Calreticulin Requires Both Its Mutant C-terminus and the Thrombopoietin Receptor for Oncogenic Transformation. Cancer Discov. 2016;6(4):368–381.

8. Venkatesan A, Geng J, Kandarpa M, et al. Mechanism of mutant calreticulin-mediated activation of the thrombopoietin receptor in cancers. J Cell Biol. 2021;220(7):e202009179.

9. Nivarthi H, Chen D, Cleary C, et al. Thrombopoietin receptor is required for the oncogenic function of CALR mutants. Leukemia. 2016;30(8):1759–1763.

10. Pecquet C, Chachoua I, Roy A, et al. Calreticulin mutants as oncogenic rogue chaperones for TpoR and traffic-defective pathogenic TpoR mutants. Blood. 2019;133(25):2669–2681.

11. Gao B, Adhikari R, Howarth M, et al. Assembly and antigen-presenting function of MHC class I molecules in cells lacking the ER chaperone calreticulin. Immunity. 2002;16(1):99–109.

12. Del Cid N, Jeffery E, Rizvi SM, et al. Modes of calreticulin recruitment to the major histocompatibility complex class I assembly pathway. J Biol Chem. 2010;285(7):4520–4535.

13. Wearsch PA, Peaper DR, Cresswell P. Essential glycan-dependent interactions optimize MHC class I peptide loading. Proc Natl Acad Sci U S A. 2011;108(12):4950–4955.

14. Desikan H, Kaur A, Pogozheva ID, Raghavan M. Effects of calreticulin mutations on cell transformation and immunity. J Cell Mol Med. 2023;27(8):1032–1044.

15. Sadasivan B, Lehner PJ, Ortmann B, Spies T, Cresswell P. Roles for calreticulin and a novel glycoprotein, tapasin, in the interaction of MHC class I molecules with TAP. Immunity. 1996;5(2):103–114.

16. Blum JS, Wearsch PA, Cresswell P. Pathways of antigen processing. Annu Rev Immunol. 2013;31:443–473.

17. Blees A, Januliene D, Hofmann T, et al. Structure of the human MHC-I peptide-loading complex. Nature. 2017;551(7681):525–528.

18. Barker DJ, Maccari G, Georgiou X, et al. The IPD-IMGT/HLA Database. Nucleic Acids Res. 2023;51(D1):D1053–D1060.

19. Bjorkman PJ, Parham P. Structure, function, and diversity of class I major histocompatibility complex molecules. Annu Rev Biochem. 1990;59:253–288.

20. Sarkizova S, Klaeger S, Le PM, et al. A large peptidome dataset improves HLA class I epitope prediction across most of the human population. Nat Biotechnol. 2020;38(2):199–209.

21. Olson E, Geng J, Raghavan M. Polymorphisms of HLA-B: influences on assembly and immunity. Curr Opin Immunol. 2020;64:137–145.

22. Zaitoua AJ, Kaur A, Raghavan M. Variations in MHC class I antigen presentation and immunopeptidome selection pathways. F1000Res. 2020;9.

23. Peh CA, Burrows SR, Barnden M, et al. HLA-B27-restricted antigen presentation in the absence of tapasin reveals polymorphism in mechanisms of HLA class I peptide loading. Immunity. 1998;8(5):531–542.

24. Zernich D, Purcell AW, Macdonald WA, et al. Natural HLA class I polymorphism controls the pathway of antigen presentation and susceptibility to viral evasion. J Exp Med. 2004;200(1):13–24.

25. Rizvi SM, Salam N, Geng J, et al. Distinct Assembly Profiles of HLA-B Molecules. The Journal of Immunology. 2014;192(11):4967–4976.

26. Geng J, Zaitouna AJ, Raghavan M. Selected HLA-B allotypes are resistant to inhibition or deficiency of the transporter associated with antigen processing (TAP). PLoS Pathog. 2018;14(7):e1007171.

27. Bashirova AA, Viard M, Naranbhai V, et al. HLA tapasin independence: broader peptide repertoire and HIV control. Proc Natl Acad Sci U S A. 2020;117(45):28232–28238.

28. Henderson RA, Michel H, Sakaguchi K, et al. HLA-A2.1-associated peptides from a mutant cell line: a second pathway of antigen presentation. Science. 1992;255(5049):1264–1266.

29. Wei ML, Cresswell P. HLA-A2 molecules in an antigen-processing mutant cell contain signal sequence-derived peptides. Nature. 1992;356(6368):443–446.

30. Arshad N, Cresswell P. Tumor-associated calreticulin variants functionally compromise the peptide loading complex and impair its recruitment of MHC-I. J Biol Chem. 2018;293(25):9555–9569.

31. Jeromin S, Kohlmann A, Meggendorfer M, et al. Next-generation deep-sequencing detects multiple clones of CALR mutations in patients with BCR-ABL1 negative MPN. Leukemia. 2016;30(4):973–976.

32. Mohan HM, Yang B, Dean NA, Raghavan M. Calreticulin enhances the secretory trafficking of a misfolded alpha-1-antitrypsin. J Biol Chem. 2020;295(49):16754–16772.

33. Garcia-Beltran WF, Holzemer A, Martrus G, et al. Open conformers of HLA-F are high-affinity ligands of the activating NK-cell receptor KIR3DS1. Nat Immunol. 2016;17(9):1067–1074.

34. Fosselteder J, Pabst G, Sconocchia T, et al. Human gene-engineered calreticulin mutant stem cells recapitulate MPN hallmarks and identify targetable vulnerabilities. Leukemia. 2023;37(4):843–853.

35. Barnstable CJ, Bodmer WF, Brown G, et al. Production of monoclonal antibodies to group A erythrocytes, HLA and other human cell surface antigens-new tools for genetic analysis. Cell. 1978;14(1):9–20.

36. Parham P, Barnstable CJ, Bodmer WF. Use of a monoclonal antibody (W6/32) in structural studies of HLA-A,B,C, antigens. J Immunol. 1979;123(1):342–349.

37. Parham P, Brodsky FM. Partial purification and some properties of BB7.2. A cytotoxic monoclonal antibody with specificity for HLA-A2 and a variant of HLA-A28. Hum Immunol. 1981;3(4):277–299.

38. Yarzabek B, Zaitouna AJ, Olson E, et al. Variations in HLA-B cell surface expression, half-life and extracellular antigen receptivity. Elife. 2018;7.

39. Kaur A, Venkatesan A, Kandarpa M, Talpaz M, Raghavan M. Lysosomal degradation targets mutant calreticulin and the thrombopoietin receptor in myeloproliferative neoplasms. Blood Adv. 2024;8(13):3372–3387.

40. El-Khoury M, Cabagnols X, Mosca M, et al. Different impact of calreticulin mutations on human hematopoiesis in myeloproliferative neoplasms. Oncogene. 2020;39(31):5323–5337.

41. Liu P, Zhao L, Loos F, et al. Immunosuppression by Mutated Calreticulin Released from Malignant Cells. Mol Cell. 2020;77(4):748–760.e749.

42. Pecquet C, Papadopoulos N, Balligand T, et al. Secreted Mutant Calreticulins As Rogue Cytokines in Myeloproliferative Neoplasms. Blood. 2023;141(8):917–929.

43. Greiner JW, Hand PH, Noguchi P, Fisher PB, Pestka S, Schlom J. Enhanced expression of surface tumor-associated antigens on human breast and colon tumor cells after recombinant human leukocyte alpha-interferon treatment. Cancer Res. 1984;44(8):3208–3214.

44. Boyer CM, Dawson DV, Neal SE, et al. Differential induction by interferons of major histocompatibility complex-encoded and non-major histocompatibility complex-encoded antigens in human breast and ovarian carcinoma cell lines. Cancer Res. 1989;49(11):2928–2934.

45. Cornel AM, Mimpen IL, Nierkens S. MHC Class I Downregulation in Cancer: Underlying Mechanisms and Potential Targets for Cancer Immunotherapy. Cancers (Basel). 2020;12(7).

46. Schurch PM, Malinovska L, Hleihil M, et al. Calreticulin mutations affect its chaperone function and perturb the glycoproteome. Cell Rep. 2022;41(8):111689.

47. Stengel A, Jeromin S, Haferlach T, Meggendorfer M, Kern W, Haferlach C. Detection and characterization of homozygosity of mutated CALR by copy neutral loss of heterozygosity in myeloproliferative neoplasms among cases with high CALR mutation loads or with progressive disease. Haematologica. 2019;104(5):e187–e190.

48. Wearsch PA, Jakob CA, Vallin A, Dwek RA, Rudd PM, Cresswell P. Major histocompatibility complex class I molecules expressed with monoglucosylated N-linked glycans bind calreticulin independently of their assembly status. J Biol Chem. 2004;279(24):25112–25121.

49. Domnick A, Winter C, Susac L, et al. Molecular basis of MHC I quality control in the peptide loading complex. Nat Commun. 2022;13(1):4701.

50. Fisette O, Schroder GF, Schafer LV. Atomistic structure and dynamics of the human MHC-I peptide-loading complex. Proc Natl Acad Sci U S A. 2020;117(34):20597–20606.

51. Frickel EM, Riek R, Jelesarov I, Helenius A, Wuthrich K, Ellgaard L. TROSY-NMR reveals interaction between ERp57 and the tip of the calreticulin P-domain. Proc Natl Acad Sci U S A. 2002;99(4):1954–1959.

52. Dick TP, Bangia N, Peaper DR, Cresswell P. Disulfide bond isomerization and the assembly of MHC class I-peptide complexes. Immunity. 2002;16(1):87–98.

53. Howe C, Garstka M, Al-Balushi M, et al. Calreticulin-dependent recycling in the early secretory pathway mediates optimal peptide loading of MHC class I molecules. EMBO J. 2009;28(23):3730–3744.

54. Cassinat B, Verger E, Kiladjian JJ. Interferon alfa therapy in CALR-mutated essential thrombocythemia. N Engl J Med. 2014;371(2):188–189.

55. Verger E, Cassinat B, Chauveau A, et al. Clinical and molecular response to interferon-alpha therapy in essential thrombocythemia patients with CALR mutations. Blood. 2015;126(24):2585–2591.

56. Hasselbalch HC, Holmstrom MO. Perspectives on interferon-alpha in the treatment of polycythemia vera and related myeloproliferative neoplasms: minimal residual disease and cure? Semin Immunopathol. 2019;41(1):5–19.

57. Skov V, Riley CH, Thomassen M, et al. The impact of interferon-alpha2 on HLA genes in patients with polycythemia vera and related neoplasms. Leuk Lymphoma. 2017;58(8):1914–1921.

58. Cimen Bozkus C, Roudko V, Finnigan JP, et al. Immune Checkpoint Blockade Enhances Shared Neoantigen-Induced T-cell Immunity Directed against Mutated Calreticulin in Myeloproliferative Neoplasms. Cancer Discov. 2019;9(9):1192–1207.

59. Rumi E, Cazzola M. Diagnosis, risk stratification, and response evaluation in classical myeloproliferative neoplasms. Blood. 2017;129(6):680–692.

60. Battle R, Poole K, Haywood-Small S, Clark B, Woodroofe MN. Molecular characterisation of the monocytic cell line THP-1 demonstrates a discrepancy with the documented HLA type. Int J Cancer. 2013;132(1):246–247.

