## Supplementary Information for "Effects of Calreticulin Mutations on HLA Class I Expression in Myeloproliferative Neoplasms"

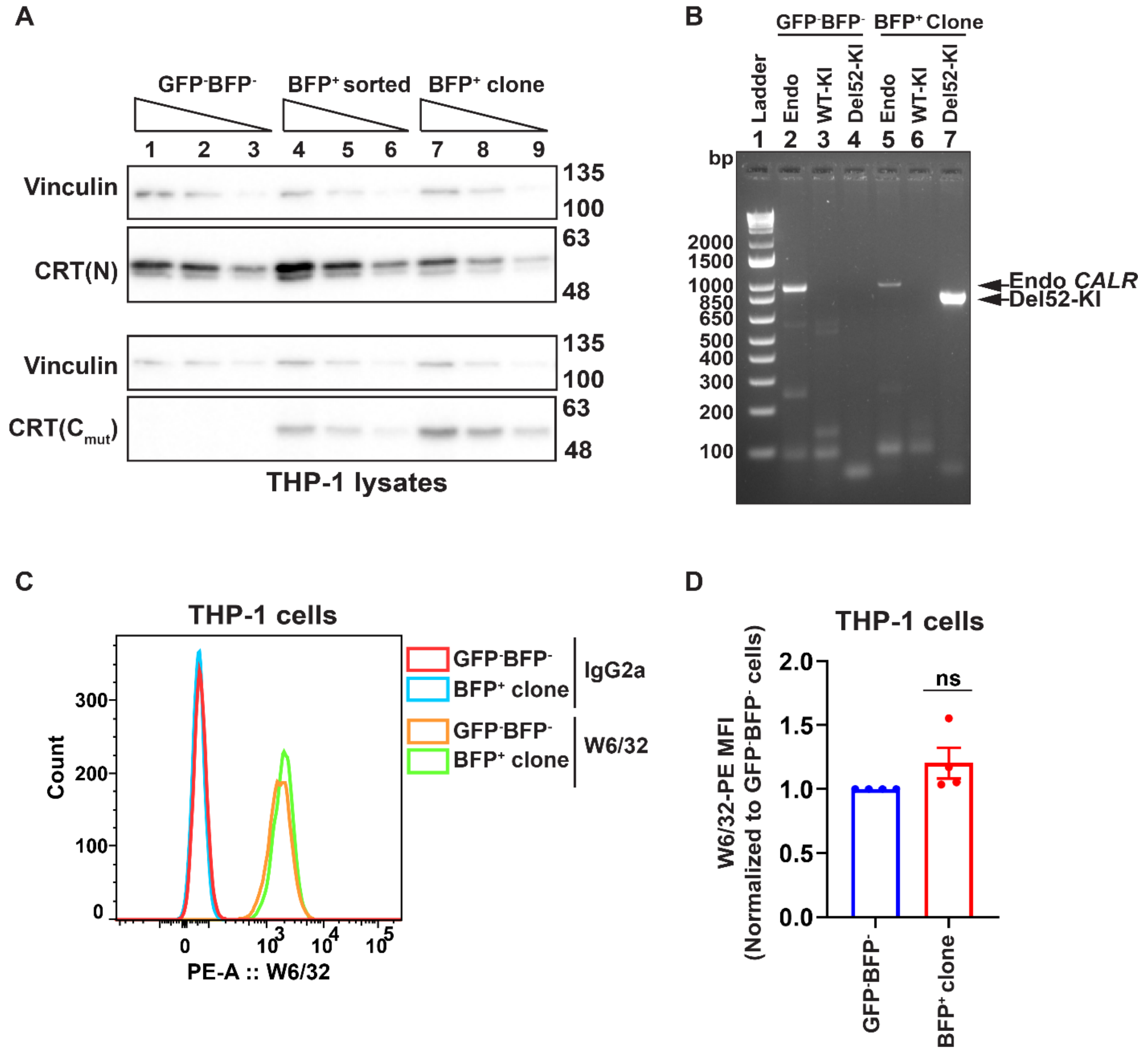

**Supplementary Figure 1: Consequences of CRT haploinsufficiency upon surface HLA class I expression in THP-1 cells. (A)** THP-1 cells were treated with Cas9/RNP and transduced with rAAV6 virus to knock in the CRT$_{Del52}$-specific exon 9 sequence at the endogenous *CALR* locus, as published earlier[1]. Representative immunoblots show wild-type CRT and mutant CRT expression detected using anti-CRT(N) and anti-CRT(C$_{mut}$) antibodies, respectively, in the lysates of sorted THP-1 GFP$^-$BFP$^-$ and BFP$^+$ cells and THP-1 BFP$^+$ single cell clone. Vinculin is shown as loading control. **(B)** Seamless integration of the *CRT$_{Del52}$*-specific knock-in fragment into the endogenous *CALR* locus in THP-1 BFP$^+$ clone was verified by in-out PCR using genomic DNA as template and compared to the GFP$^-$BFP$^-$ THP-1 cells. Endogenous *CALR* (Endo), Del52 knock-in (Del52-KI) and wild-type knock-in (WT-KI) fragments were amplified using a common forward primer and specific reverse primers. The agarose gel image shows PCR products amplified from endogenous *CALR* (Endo *CALR*) locus (~919 bp) and *CRT$_{Del52}$* knock-in (Del52-KI) allele (~746 bp) as indicated. **(C and D)** Representative histograms show surface staining with W6/32-PE or

Isotype control (IgG2a-PE) antibody measured by flow cytometry on GFP$^-$BFP$^-$ THP-1 cells or BFP$^+$ clone (C). The graph shows the average MFI value of surface staining with W6/32 antibody in THP-1 BFP$^+$ clone normalized to the MFI values for GFP$^-$BFP$^-$ THP-1 cells, calculated from four independent experiments (D). The graph was plotted using GraphPad Prism, and statistical significance was calculated using a one-sample t-test.

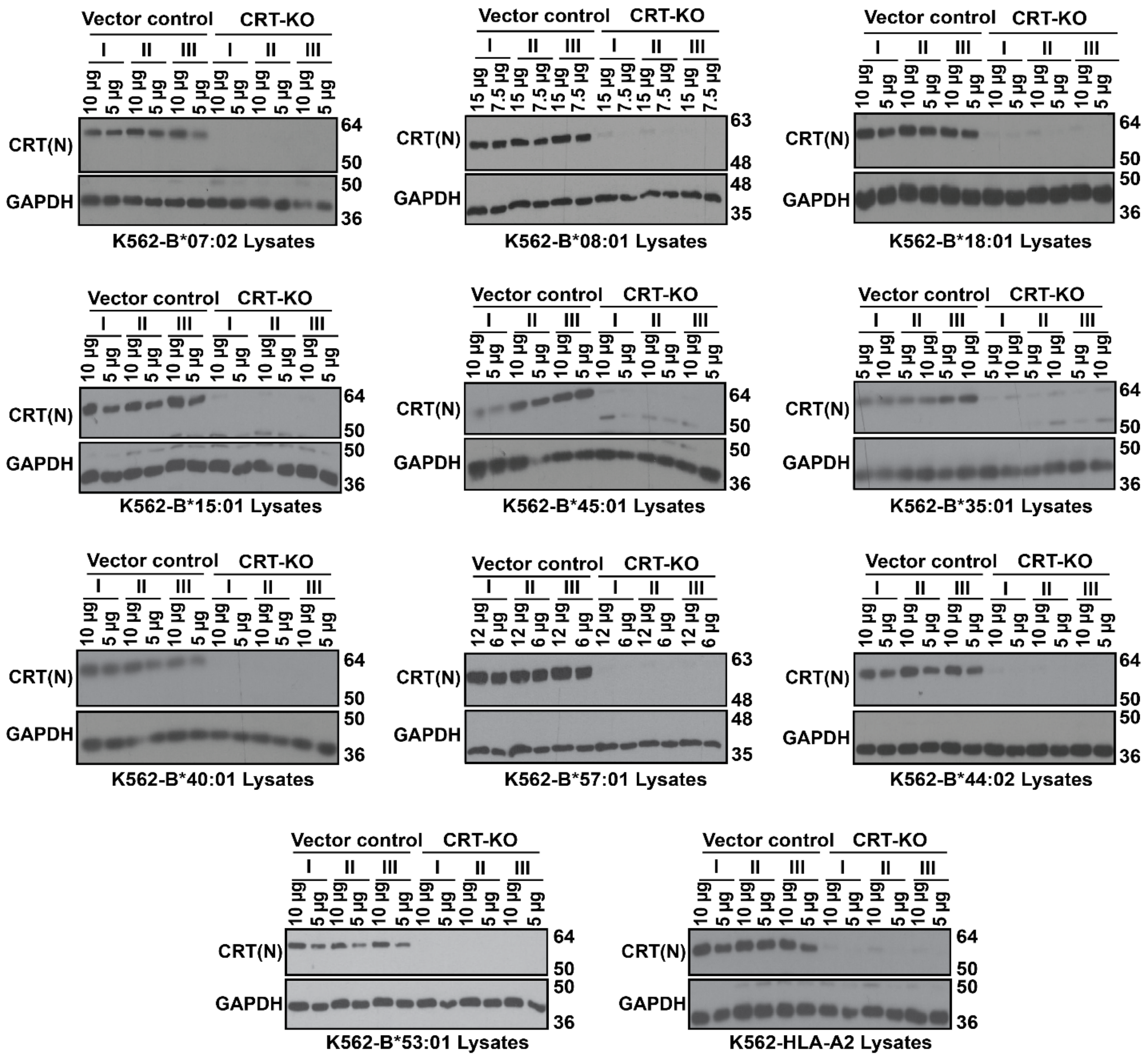

**Supplementary Figure 2: CRISPR/Cas9-based CRT knockdown in K562 cells:** HLA class I null K562 cells were transduced with retroviruses to express individual HLA class I allotypes (monoallelic) and were then subjected to CRISPR/Cas9-based knockdown of CRT expression (CRT-KO). Representative blots show CRT protein levels in K562 CRT-KO cells-expressing individual HLA class I allotypes compared to cells transduced with empty vector (Vector Control). Two different amounts of lysates from each of the three biological replicates (I, II and III) of CRT-KO and vector control cells were loaded as indicated. CRT was probed using anti-CRT(N) antibody and GAPDH is shown as a loading control.

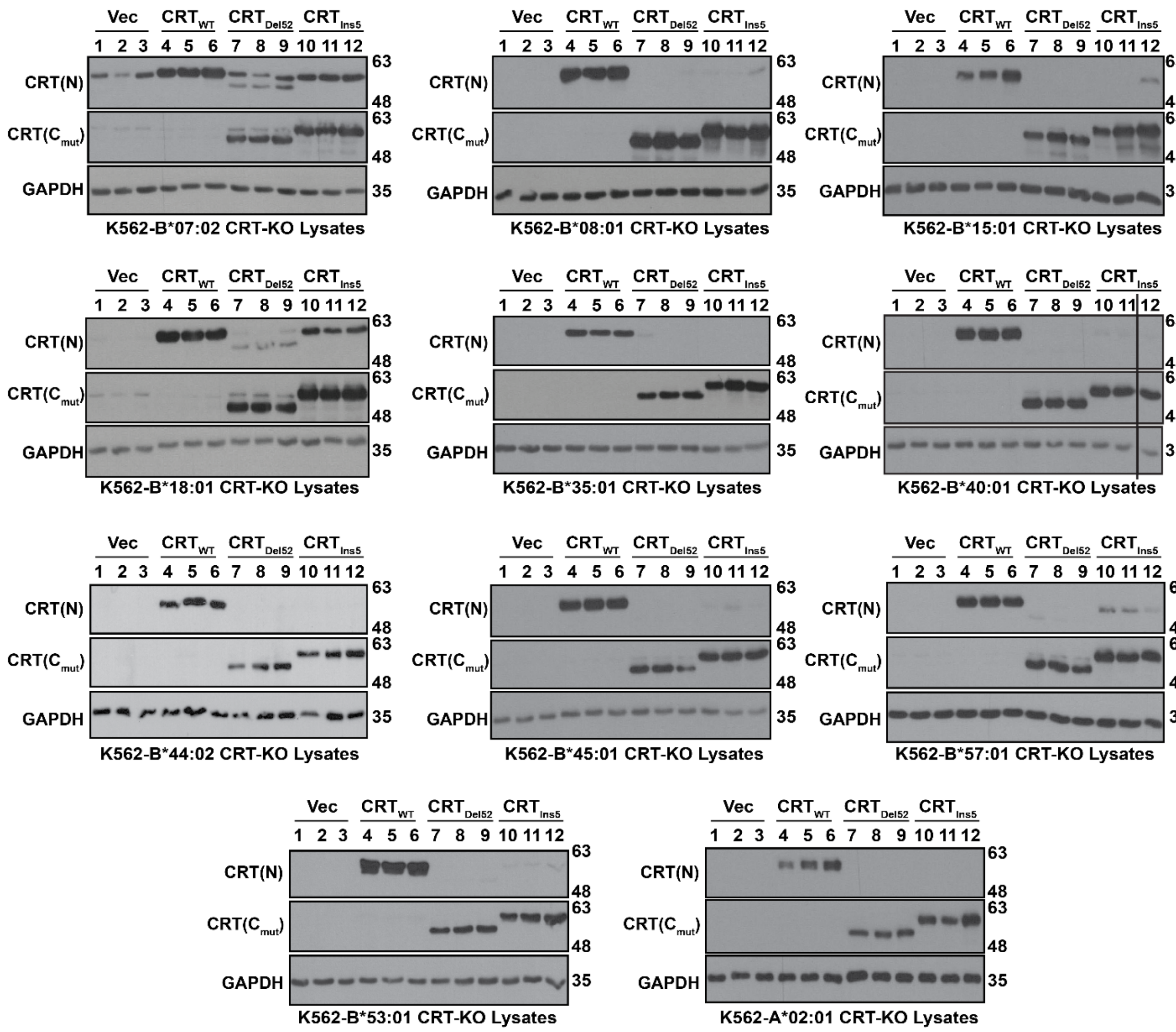

**Supplementary Figure 3: Reconstitution of expression of wild-type CRT or MPN-linked CRT mutants in K562-CRT-KO cells:** HLA class I null K562 CRT-KO cells-expressing individual HLA class I allotypes as shown were transduced with retrovirus to reconstitute either wild-type CRT (CRT$_{WT}$) or MPN-linked CRT mutant (CRT$_{Del52}$ or CRT$_{Ins5}$) expression. Representative blots show wild-type or mutant CRT protein levels detected using anti-CRT(N) or anti-CRT(C$_{mut}$) antibody, respectively, in K562 CRT-KO cells, following retroviral transduction-based reconstitution of the expression of indicated CRT protein. Lysates from three biological replicates of retroviral transduction with either empty vector (Vec), wild-type CRT (CRT$_{WT}$) or mutants (CRT$_{Del52}$ or CRT$_{Ins5}$) were loaded. GAPDH is shown as a loading control.

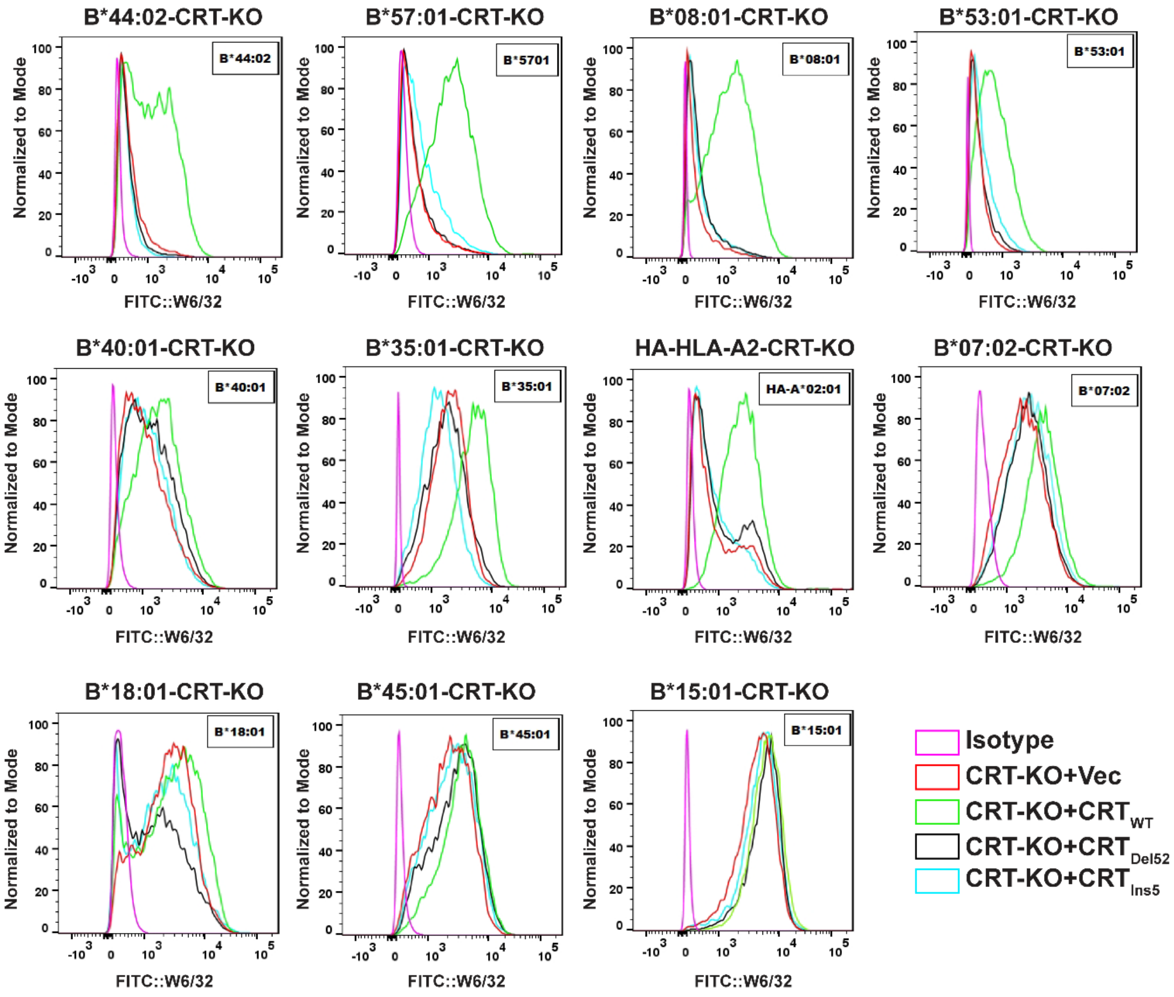

**Supplementary Figure 4: Reconstitution of wild-type CRT expression, but not the MPN-linked CRT mutants, rescues surface HLA class I levels in CRT-KO cells.** Flow cytometry of K562 CRT-KO cells expressing indicated HLA class I allotypes and reconstituted with either wild-type CRT (CRT-KO+CRT$_{WT}$) or mutant CRT (CRT-KO+CRT$_{Del52}$ or CRT-KO+CRT$_{Ins5}$) proteins. K562 CRT-KO cells transduced with empty vector (CRT-KO+Vec) are included as a control. Representative histograms show surface staining with the W6/32 antibody or the isotype control antibody.

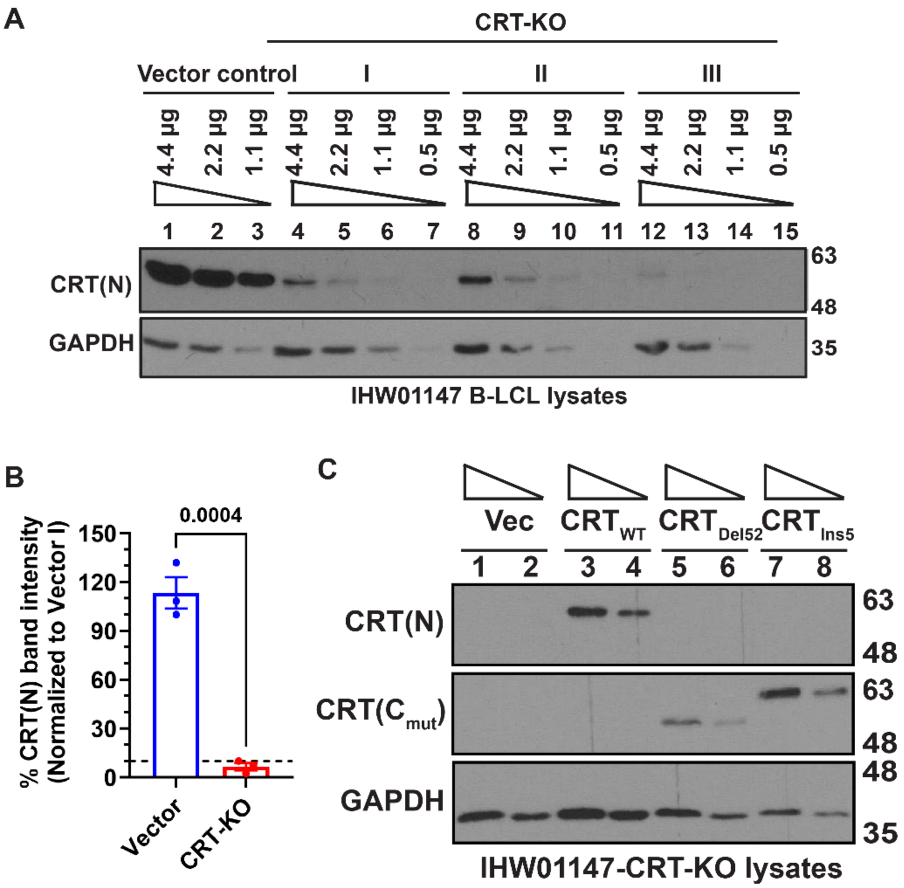

**Supplementary Figure 5: CRISPR/Cas9-based CRT knockdown and reconstitution of mutant CRT expression in IHW01147 B-LCL. (A)** Representative blots show CRT protein levels in IHW01147 B-LCL transduced with lentivirus packaged with either empty vector (Vector control) or the vector containing *CALR*-targeting sgRNA for CRISPR/Cas9-based knockdown of CRT expression (CRT-KO). Three different lysate loads from vector control cells and each of the three biological replicates (I, II and III) of CRT-KO were loaded as indicated. **(B)** Quantification of CRT protein levels from immunoblots of lysates of IHW01147 CRT-KO cells compared to vector control cells. Image J was used to quantify band intensities on immunoblots. The CRT intensities were normalized to the GAPDH intensities. The average percent CRT intensities calculated from the values of three vector-control and CRT-KO cell lines, assuming CRT intensity for the vector I cell line as 100%, are plotted. The statistical significance was calculated using an unpaired-t test in GraphPad Prism. **(C)** Representative blots show wild-type and mutant CRT expression in the lysates of IHW01147-CRT-KO cells following reconstitution with either wild-type CRT ($CRT_{WT}$) or mutant ($CRT_{Del52}$ or $CRT_{Ins5}$) proteins. IHW01147-CRT-KO cells transduced with empty vector (Vec) are also included. In panels A and C, wild-type and mutant CRT proteins were detected using anti-CRT(N) and anti-CRT($C_{mut}$) antibodies, respectively, and GAPDH is shown as a loading control.

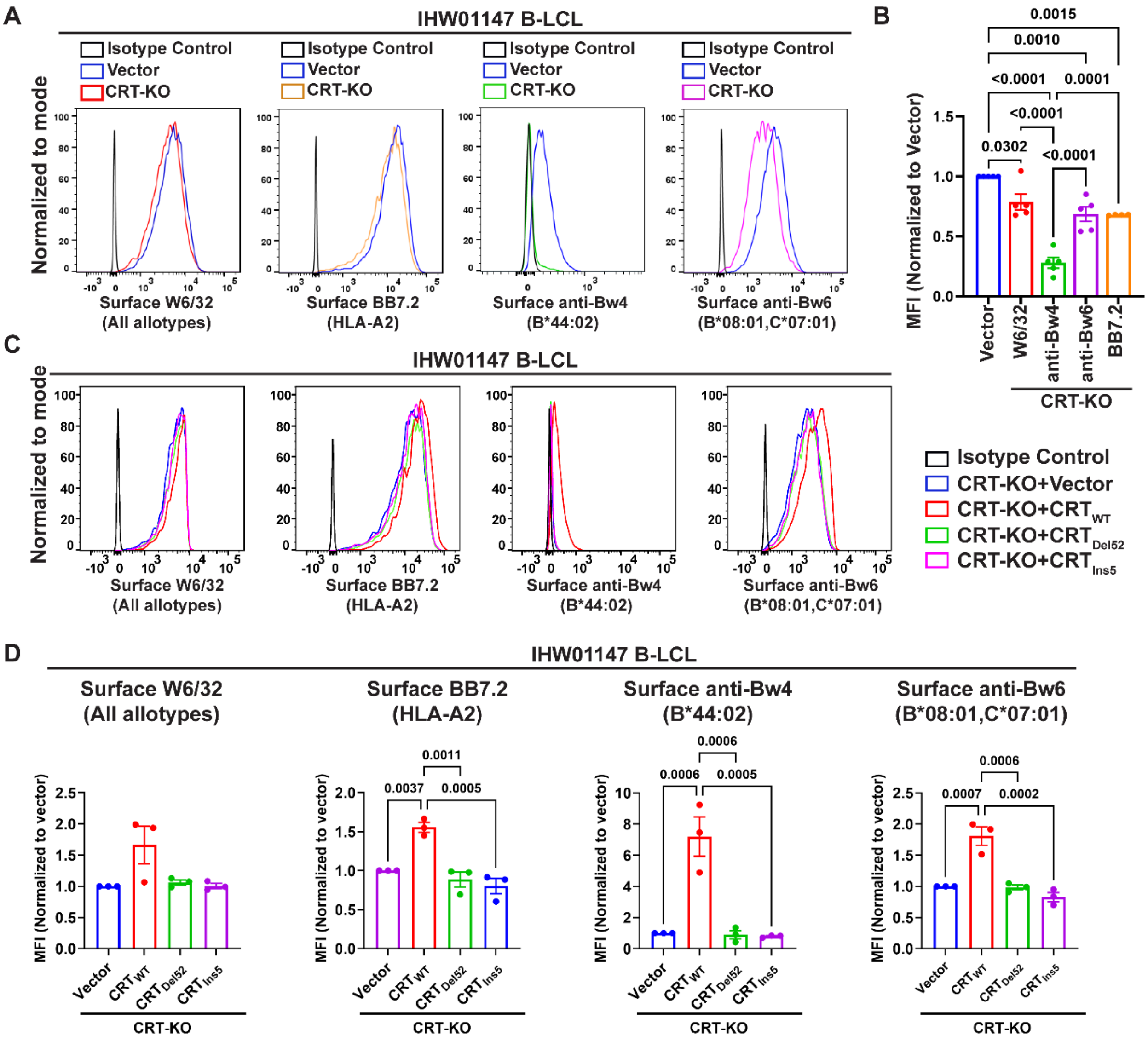

**Supplementary Figure 6: Consequences of full calreticulin deficiency and MPN calreticulin reconstitution on HLA class I expression in IHW01147 B-LCL. (A and B)** Surface HLA class I expression was measured by flow cytometry using the W6/32, anti-Bw4, anti-Bw6, or BB7.2 antibodies in the IHW01147 cells (A*01:01/02:01, B*08:01/44:02, C*07:01/05:01) following CRT knockdown and compared to the vector control cells. Representative histograms show surface staining of IHW01147-CRT-KO cells with the indicated anti-HLA class I antibody or the isotype control antibody (A), and the bar graph shows the averaged data (B). Each dot shows the average MFI value calculated from three biological replicates of CRT knockdown cells, stained in parallel across multiple independent experiments (n=5 for W6/32, anti-Bw4, anti-Bw6, and n=4 for BB7.2). **(C and D)** The IHW01147-CRT-KO B-LCL reconstituted with wild-type CRT (CRT-KO+CRT$_{WT}$) or MPN-linked CRT mutants (CRT-KO+CRT$_{Del52}$ or CRT-KO+CRT$_{Ins5}$) or the vector control cells (CRT-KO+Vector) were stained with the isotype control or indicated anti-HLA class I antibody. Representative histograms show surface HLA class I expression measured with the indicated antibodies (C), and graphs show averaged data (D) from independent flow cytometry experiments (n=3). Statistical significance in panels B and D was calculated in GraphPad Prism using ordinary one-way ANOVA.

| S.No | Healthy donor/ Patient ID | Mutation | Clinical Treatment | Disease, Lab VAF, or clinical VAF | Classical Monocyte Fraction among live PBMC | W6/32 ABC platelets | W6/32 ABC Classical Monocytes | Classical Monocyte Anti-CRT (N) Ratios | Classical Monocyte Anti-CRT($C_{mut}$) Ratios |
|---|---|---|---|---|---|---|---|---|---|
| 1 | ***CALR*7438** | *CALR* (Ins5) | Pegylated IFN | ET 28.56 | 2.926 | | 2374926.554 | 2.489 | 1.611 |
| 2 | ***CALR*8178(A)** ***CALR*8178(B)** | *CALR* (Ins5) | Hydroxyurea & Pegylated IFN | ET 9.02 (B) | 2.275 (A) 2.334 (B) | 40958.079 (B) | 1915640.437 (A) 2995756.220 (B) | 2.839 (A) 2.406 (B) | 0.936 (A) 1.987 (B) |
| 3 | ***CALR*4401** | *CALR* (Ins5) | Pegylated IFN | ET 4.69 | 4.855 | 92288.011 | 3547358.41 | 2.307 | 1.988 |
| 4 | ***CALR*4725** | *CALR* (Del52) | Pegylated IFN & Aspirin | ET 1.13 | 7.832 | 34677.659 | 1463075.204 | 2.193 | 1.511 |
| 5 | ***CALR*9845** | *CALR* (Ins5) | Hydroxyurea & Xarelto | ET 29.13 | 13.104 | | 1846640.970 | 1.970 | 1.502 |
| 6 | ***CALR*7557** | *CALR* (Del52) | Hydroxyurea | ET 0.72 | 8.023 | | 2150330.301 | 1.560 | 0.837 |
| 7 | ***CALR*2288** | *CALR* (Del52) | Hydroxyurea & Aspirin | ET 31.2# | | 26777.506 | | | |
| 8 | ***CALR*7471** | *CALR* (Ins5) | Hydroxyurea & Plavix | Pre-PMF 76.2# | 8.982 | 15121.281 | 872521.439 | 2.217 | 1.205 |
| 9 | ***CALR*8251** | *CALR* (Ins5) | Ruxolitinib & Hydroxyurea | Post ET-MF 91.3 | 6.018 | 17539.712 | 896655.371 | 11.604 | 1.850 |
| 10 | ***CALR*7610** | *CALR* (Ins5) | Ruxolitinib | Post ET-MF 21.45 | 1.944 | | 886639.456 | 2.641 | 1.420 |
| 11 | ***CALR*9342** | *CALR* (Del52) | Ruxolitinib | Post ET-MF 33.9# | 0.791 | 22624.090 | 1450642.144 | 4.048 | 2.299 |
| 12 | ***CALR*1983** | *CALR* (Del34) | Ruxolitinib | Post ET-MF 31.3 | 3.559 | 22312.841 | 628808.804 | 1.965 | 0.686 |
| 13 | ***CALR*0266** | *CALR* (Del52) | LSD1 inhibitor | Post ET-MF 1.22 | 16.841 | 51689.279 | 1545874.332 | | |
| 14 | ***CALR*0387** | *CALR* (Del52) | Danazol | PMF 1.62 | 0.435 | 61213.889 | 1169475.406 | 5.150 | 3.369 |
| 15 | ***CALR*2077** | *CALR* (Del52) | PRBC transfusion | PMF 0.82 | 0.948 | 27455.839 | 1439724.413 | 4.675 | 2.951 |
| 16 | ***CALR*5628** | *CALR* (Del52) | Aspirin | PMF 13.5# | | 23087.115 | | | |

| | | | | | | | |
|---|---|---|---|---|---|---|---|
| 17 | ***JAK*6440** | *JAK*2 V617F | Aspirin & Ruxolitinib | PV | 9.734 | 28881.544 | 692761.752 |
| 18 | ***JAK*0621** | *JAK*2 V617F | Hydroxyurea & Aspirin | PV | 1.637 | 17759.310 | ***482450.924*¶** |
| 19 | ***JAK*0196** | *JAK*2 V617F | Hydroxyurea & Aspirin | PV | 6.122 | 43247.766 | 1713080.441 |
| 20 | ***JAK*1761** | *JAK*2 V617F | Aspirin | PV | 1.284 | 16035.165 | 2045781.240 |
| 21 | ***JAK*5092** | *JAK*2 V617F | Fedratinib | Post-PV MF | 0.088 | 23010.722 | ***2032950.519*¶** |
| 22 | ***JAK*0922** | *JAK*2 V617F | Ruxolitinib | PMF | 4.049 | 11963.067 | 1143270.398 |
| 23 | ***MPL*8524** | *MPL* W515K | BET inhibitor 2018-090 | PMF | 0.319 | 63881.640 | 1032644.214 |
| 24 | **HD1** | | | | 9.425 | | 1555580.447 |
| 25 | **HD2** | | | | 19.520 | | 1336275.326 |
| 26 | **HD4** | | | | 5.386 | | 1052015.857 |
| 27 | **HD5** | | | | 4.301 | | 1478268.031 |
| 28 | **HD6** | | | | 23.166 | 63485.073 | 1186217.548 |
| 29 | **HD7** | | | | 9.048 | 83400.905 | 1876420.004 |
| 30 | **HD8** | | | | 6.689 | 56722.238 | 1854469.143 |
| 31 | **HD9(A)**<br>**HD9(B)** | | | | 1.421<br>3.540 | 30797.939 (A)<br>73674.249 (B) | 1576273.352 (A)<br>1568532.596 (B) |
| 32 | **HD10(A)**<br>**HD10(B)**<br>**HD10(C)**<br>**HD10(D)** | | | | 1.284 (A)<br>9.923 (B)<br>9.570 (C)<br>6.373 (D) | 34666.741 (A)<br>23119.198 (B)<br>40634.265 (C)<br>38199.731 (D) | ***1293488.098*¶** (A)<br>1517286.667 (B)<br>863553.733 (C)<br>1399483.013 (D) |
| 33 | **HD11(A)**<br>**HD11(B)** | | | | 7.863 (A)<br>5.579 (B) | 52911.981 (A) | 1527245.391 (A)<br>2943547.538 (B) |
| 34 | **HD12** | | | | 2.611 | 46650.580 | 1154325.872 |
| 35 | **HD13** | | | | 5.051 | 38754.923 | 1173267.481 |
| 36 | **HD14** | | | | 1.307 | 24285.677 | 1116187.333 |
| 37 | **HD15** | | | | 12.878 | 22905.313 | 2164293.696 |

**Supplementary Table 1: Disease, mutation type, and clinical treatment for de-identified patients and results from experimental measurements of their platelet or PBMC samples**

The table provides details of MPN patients with *CALR*, *JAK2* or *MPL* mutations and healthy donors. IDs for de-identified samples and information about the driver mutation, disease, and treatment was obtained from the Myeloproliferative Disease Repository (study ID: HUM0006778). The *CALR* variant allele frequency (VAF) values were measured using genomic DNA isolated from PBMC or obtained for some patient samples as part of the clinical information (indicated by #). VAF values were measured in the lab using the same PBMC samples as those used for surface HLA class I measurements, except for *CALR*8251, for which the VAF was measured with a blood sample collected six years and 11 months prior to the sample collection for surface HLA class I measurements. Clinical VAF values were measured using peripheral blood 1 year 8 months before surface HLA measurements in *CALR*2288 PBMC, 3 months prior to surface HLA measurements in *CALR*7471 PBMC, 4 years 5 months before surface HLA measurements in *CALR*5628 PBMC, and 1 year 8 months after surface HLA class I measurements in *CALR*9342 PBMC. Fractions of classical monocytes among live PBMC for patient samples and healthy donors, and ABC values derived from surface staining of platelets and classical monocytes with the W6/32 antibody are also shown. The MFI values for surface staining with anti-CRT(N) and anti-CRT($C_{mut}$) antibodies of classical monocytes from MPN patients with *CALR* mutations are provided as a ratio relative to the corresponding MFI values of classical monocytes derived from the same-day healthy donors. Multiple collections from same patient or healthy donor are indicated by an alphabet enclosed within parentheses added to the ID, VAF value, ABC values or anti-CRT MFI ratios. The ABC values italicized and bold-faced (indicated by ¶) were excluded because the number of events within the specific gate was <100. Patients determined to have a prevalence of cells with homozygous *CALR* mutations are highlighted in red color, while those with high PBMC VAF but showing expression of both mutant and wild-type CRT in platelet lysates, consistent with the prevalence of cells with heterozygous *CALR* mutations, are highlighted in yellow color. Healthy donor sample HD3 and *CALR*1596 patient sample (PMF patient with Del52 mutation prescribed Ruxolitinib treatment) were included in the immunoblots of platelet lysates, but not in the surface HLA class I expression analyses. BET, Bromodomain and extraterminal protein; IFN, Interferon; PRBC, Packed Red Blood Cell; LSD1, Lysine-specific demethylase 1

| S. No | Sample ID | Healthy/MPN | HLA-A | | HLA-B | | HLA-C | |
|---|---|---|---|---|---|---|---|---|
| 1 | ***CALR*7471** | CALR (Ins5) | A*11:01 | A*29:02 | B*14:02 | B*44:02 | C*08:02 | C*05:01 |
| 2 | ***CALR*8251** | CALR (Ins5) | A*24:02 | A*25:01 | B*08:01 | B*18:01 | C*07:01 | C*12:03 |
| **HLA-A2⁺ MPN patients and healthy donors** | | | | | | | | |
| 1 | **HD5** | Healthy | ***A*02:01*** | A*24:02 | B*44:02 | B*57:02 | C*05:01 | C*18:02 |
| 2 | **HD12** | Healthy | ***A*02:01*** | A*32:01 | B*53:01 | B*27:05 | C*04:01 | C*01:02 |
| 3 | **HD13** | Healthy | ***A*02:01*** | A*01:01 | B*15:01 | B*08:01 | C*03:03 | C*07:01 |
| 4 | **HD14** | Healthy | ***A*02:01*** | A*24:02 | B*51:01 | B*35:01 | C*15:02 | C*04:04 |
| 5 | **HD15** | Healthy | ***A*02:01*** | A*03:01 | B*51:01 | B*07:02 | C*15:02 | C*07:02 |
| 6 | ***CALR*0266** | *CALR* (Del52) | ***A*02:01*** | A*03:01 | B*44:02 | B*44:03 | C*07:04 | C*02:02 |
| 7 | ***CALR*2077** | *CALR* (Del52) | ***A*02:01*** | A*03:01 | B*27:05 | B*07:02 | C*02:02 | C*07:02 |
| 8 | ***JAK*0922*** | *JAK2*V617F | ***A*02:01*** | ***A*02:01*** | B*44:03 | B*13:02 | C*16:01 | C*06:02 |
| 9 | ***JAK*1761** | *JAK2*V617F | ***A*02:01*** | A*11:01 | B*51:01 | B*41:02 | C*02:02 | C*17:03 |

**Supplementary Table 2: HLA class I genotypes of *CALR*7471 and *CALR*8251, and the healthy donors and MPN patients used for BB7.2 ABC measurements**. The BB7.2 ABC values for sample *JAK*0922 (marked by a *) homozygous for HLA-A*02:01, were divided by 2 to use them for comparisons.

| S.No | Name | Sequence (5' to 3') | Used for | References |
|---|---|---|---|---|
| 1 | *CALR* sgRNA | GGCCACAGATGTCGGGACCT | CRISPR/Cas 9 based CRT-knockout in K562 cells | Previously published[2] |
| 2 | pLB-*CALR* sgRNA | Forward (**BsmBI**): 5'-**CACCG**GGCCACAGATGTCGGGACCT-3' | Cloning of *CALR*-sgRNA in pLentiCRISPRv2-BLAST (pLB) vector. The underlined bases correspond to the adaptor sequences for cloning. | Previously published[2] |
| | | Reverse (**BsmBI**): 5-**AAAC**AGGTCCCGACATCTGTGGCC**C**-3' | | |
| 3 | pSIP Vector minus *ZsGreen* | FP (**PmeI**)- AATTCccg**GTTTAAAC**gccG | Replacement of the *ZsGreen* fragment of the pSIP-*ZsGreen* vector with the PmeI sequence | This study |
| | | RP (**PmeI**)- Gggc**CAAATTTG**cggCCTAG | | |
| 4 | pSIP-$CRT_{WT}$ | FP (**EcoRI**)- CGAA**GAATTC**GCCGCCACCATGCTGCTATCCG | Cloning of $CRT_{WT}$ cDNA in pSIP vector | This study |
| | | RP (**NotI**)- GCATTATT**GCGGCCGC**CTACAGCTCGTCCTTGGC | | |
| 5 | pSIP-$CRT_{Del52}$ or $CRT_{Ins5}$ | FP (**EcoRI**)- CGAA**GAATTC**GCCGCCACCATGCTGCTATCCG | Cloning of $CRT_{Del52}$ or $CRT_{Ins5}$ cDNA in pSIP vector | This study |
| | | RP (**NotI**)- GCATTATT**GCGGCCGC**TCAGGCCTCAGTCCAGCC | | |
| 6 | pSIP-*HLA-B*15:01* | FP (**EcoRI**)- CGAA**GAATTC**GCCGCCACCATGCGGGTCACGG | Cloning of *HLA-B*15:01* into pSIP vector | This study |
| | | RP (**BamHI**)- CCAGCCG**GGATCC**GATCAAGCTGTGAGAGAC | | |
| 7 | pSIP-*HLA-B*07:02*; pSIP-*HLA-B*08:01* | FP (**EcoRI**)- CGAA**GAATTC**GCCGCCACCATGCTGGTCATGG | Cloning of *HLA-B*08:01* and *HLA-B*07:02* into pSIP vector | This study |
| | | RP (**BamHI**)- CCAGCCG**GGATCC**GATCAAGCTGTGAGAGAC | | |
| 8 | Site-directed mutagenesis (SDM) of pSIP-$CRT_{WT}$, $CRT_{Del52}$ or $CRT_{Ins5}$ | C1QF-ATACAACATCATGTTTGGGCCTGATATCTGTGGCCCTGGCACC | SDM to make pSIP-$CRT_{WT}$, $CRT_{Del52}$ or $CRT_{Ins5}$ constructs gRNA resistant | This study |
| | | C1QR-GGTGCCAGGGCCACAGATATCAGGCCCAAACATGATGTTGTAT | | |
| | | C2QF-CAACATCATGTTTGGGCCTGATATATGCGGACCTGGCACCAAGA | | |
| | | C2QR-TCTTGGTGCCAGGTCCGCATATATCAGGCCCAAACATGATGTTG | | |

| | | | | |
|---|---|---|---|---|
| 9 | gRNA for CRISPR knock-in | CGCCTGTAATCCTCGCCCAG | CRISPR Knock-in after complexing with Cas9 to make Cas9-RNP | Previously published[3] |
| 10 | Primers for PCR-based confirmation of knock-in | Common FP- AAGTGATCCGTTCGCCATGACCTCCC | Verification of the knock-in of wild-type *CALR* or $CRT_{Del52}$-specific exon 8,9 sequences at the endogenous *CALR* locus | This study |
| | | Endo RP- CTACAGCTCGTCCTTGGCCTGGCCG | | |
| | | WT-KI RP- CTAGAGTTCGTCTTTTGCTTGCCCGGGTAC | | |
| | | Del52-KI RP- CGACAGCTAGTTCTGGGCCTCGCCGG | | |
| 11 | Primers for RT-qPCR | FP- GAAGACAAGAAACGCAAAGAGG<br>RP- GGACATCTTCCTCCTCATCTTC | RT-qPCR-based measurement of wild-type *CALR* mRNA expression | This study |
| 12 | Primers for mutant *CALR* fragment amplification | FP- GGCAAGGCCCTGAGGTGT<br>RP- GGCCTCAGTCCAGCCCTG | Mutant CALR fragments were amplified using genomic DNA from patient PBMC as template and submitted to Genewiz for variant allele frequency (VAF) analysis | Previously published[4] |

**Supplementary Table 3: Sequences of primers and sgRNAs.** The restriction enzyme sites are boldfaced, and the underlined sequences are Kozak sequences. FP, Forward Primer; RP, Reverse Primer

### Supplementary Materials and Methods

#### Materials

All laboratory chemicals were purchased from Sigma. Vacutainer® ACD solution A tubes (364606) were obtained from BD. Ficoll® Paque-Plus density gradient media, apyrase, ACD solution, and protease inhibitor cocktail (P8340) were obtained from Sigma. FlowPRA single antigen HLA beads, anti-Bw6 FITC, and anti-Bw4 FITC were purchased from One Lambda. The 7-AAD dye was obtained from BD. All cell culture media, FBS, antibiotic-antimycotic, puromycin, geneticin, Puromycin, Blasticidin, Opti-MEM™ I reduced serum media, Recovery™ Cell Culture Freezing Medium, and Sodium Pyruvate were obtained from Gibco (Invitrogen). BCA assay protein estimation kit and Pierce™ ECL western blotting substrate (32106) were obtained from Pierce. QuikChange II XL Site-Directed Mutagenesis Kit was procured from Agilent. The BsmBI enzyme was purchased from Thermo Fisher Scientific. Restriction enzymes and T4 DNA ligase were purchased from New England Biolabs. Wizard® Plus SV Miniprep kit, GoTaq® Green Master Mix, and FuGENE® HD transfection reagent were purchased from Promega. Lipofectamine™ LTX with Plus™ reagent was procured from Invitrogen. The gel extraction kit, PCR purification kit, DNeasy Blood and Tissue kit, and RNeasy Plus Mini kit were procured from Qiagen.

#### Cell lines and antibiotics

K562 cells were maintained in RPMI media with 10% FBS, L-Glutamine, and antibiotic-antimycotic. Wherever indicated, blasticidin, puromycin, or geneticin drug was added to select for transduced K562 cells at 5 µg/mL, 0.5 µg/mL, and 500 µg/mL concentrations, respectively. THP-1 cells were maintained in RPMI media (ATCC modification, Gibco catalog #A10491) supplemented with 10% FBS, L-Glutamine, and antibiotic-antimycotic. Following transductions, THP-1 cells were selected using 5 µg/mL blasticidin or 2 µg/mL puromycin. The B-lymphoblastoid cell lines (B-LCLs) were maintained in RPMI media supplemented with 15% FBS, 1 mM Sodium Pyruvate, and antibiotic-antimycotic. For the drug selection of B-LCLs after transduction, 2.5 µg/mL blasticidin and 0.25 µg/mL puromycin were used wherever indicated.

#### sgRNA and gene constructs

The single guide RNA (sgRNA) targeting a sequence 5'-GGCCACAGATGTCGGGACCT-3' located at the boundary of intron 3 and exon 4 of the *CALR* gene was used for knockdown of CRT expression[2]. The oligos were procured from IDT after adding the specific adaptor sequences (underlined and boldfaced in Supplementary Table 3) for cloning at the BsmBI site of pLentiCRISPRv2-blasticidin (pLB) vector. The gRNA used for CRISPR knock-in experiments (Supplementary table 3) targeting intron 7 of the *CALR* gene was purchased from Synthego with chemical modifications (2'-O-Methyl group at 3 first and last bases and 3' phosphorothioate bonds between the first 3 and last 2 bases).

The pMSCV constructs of wild-type human calreticulin, as well as the $CRT_{Del52}$ and $CRT_{Ins5}$ mutants have been described earlier[5]. Site-directed mutagenesis (QuikChange II XL Site-Directed Mutagenesis Kit, Agilent) was used to introduce silent mutations within the sgRNA-binding region of wild-type and mutant CRT constructs using primers given in the supplementary Table 3. Six silent mutations were introduced in two rounds of mutagenesis using primers C1QF and C1QR in the first round and primers C2QF and C2QR and mutated products from round 1 as templates in the second round of mutagenesis. The PCR was set up following the manufacturer's protocol, followed by treatment with the Dpn1 enzyme for 1h at 37°C. XL10-Gold Ultracompetent bacterial cells (Agilent) were transformed with the DpnI-digested PCR product following the manufacturer's recommendations.

For reconstitution of CRT expression in CRT-KO cells, the sgRNA-resistant versions of human wild-type ($CRT_{WT}$) and mutant ($CRT_{Del52}$ and $CRT_{Ins5}$) CRT, generated as mentioned above, were cloned in the pSIP-*ZsGreen* (pLVX-SFFV-IRES-Puro)[6,7] vector. The *CALR* genes were amplified by PCR using primers given in supplementary Table 3 and GoTaq® Green Master Mix (Promega). The amplicons and the pSIP-*ZsGreen* (pLVX-SFFV-IRES-Puro) vector were digested with EcoRI and NotI enzymes. The digested vector was subjected to agarose gel electrophoresis and purification using the Qiagen gel extraction kit. The digested CRT inserts were purified using by Qiagen PCR purification kit. The insert and vector were combined in a molar ratio of 3:1 for ligation using the T4DNA ligase enzyme. The STBL3 competent *E. coli* cells were transformed with the ligation mix, and colonies were screened for the presence of the insert and verified by Sanger Sequencing.

The pMSCV neo constructs of HLA class I alleles used for expression of HLA class I in K562 cells have been described earlier[8]. The HLA-B*07:02, HLA-B*08:01, and HLA-B*15:01 were amplified using primers given in the supplementary table 3 and subcloned at the EcoRI and BamHI sites in the pSIP-*ZsGreen* (pLVX-SFFV-IRES-Puro) vector.

Digestion of the pSIP-*ZsGreen* vector with EcoRI and NotI/BamHI enzymes during the cloning of different *CALR* or HLA class I constructs allowed the removal of the *ZsGreen* fragment. The empty pSIP vector without *ZsGreen* was generated by replacing the *ZsGreen* sequence with a PmeI restriction enzyme site using primers given in supplementary table 3. The pSIP-*ZsGreen* vector was digested with EcoRI and BamHI enzymes and ligated to annealed and phosphorylated oligos, followed by transformation of *E. coli* STBL3 cells with the ligation mix.

**Construction of homology-dependent repair (HDR) templates for CRISPR knock-in**

The HDR templates used for CRISPR knock-in were designed to insert the exon 8 sequence of wild-type *CALR* fused to either wild-type exon 9 or $CRT_{Del52}$ exon 9 sequences at the cut site, as described in a recent study[3]. The left homology arm (LHA) consisted of 242 bases of *CALR* sequence upstream of the cut site for Cas9 enzyme within the gRNA target sequence, and the right homology arm (RHA) extended 400 bases downstream from the cut site. The splice acceptor arm included a sequence identical to the 150 bases upstream of the exon 8 in the wild-type *CALR* gene and was placed after the LHA within the HDR templates. Exon 8 and 9 sequences in the HDR templates were modified compared to the endogenous *CALR* exon 8 and 9 sequences using codon optimization. The wild-type *CALR*-specific knock-in template included a GFP reporter gene, and the $CRT_{Del52}$-specific knock-in template included a BFP reporter gene. The specific reporter gene was added downstream of the *CALR* knock-in fragment sequence and the spleen focus-forming virus (SFFV) promoter. An SV40 polyadenylation sequence was added after the wild-type or mutant exon 9 sequence, and a bovine growth hormone (bGH) polyadenylation sequence was added after the reporter gene sequence. The HDR templates were synthesized by GenScript and subcloned at the BamHI and NotI sites of pAAV-MCS2 vector (Addgene, Plasmid #46954).

**AAV generation**

Recombinant adeno-associated virus (AAV) packaged with the HDR templates carrying wild-type *CALR* or $CRT_{Del52}$-specific knock-in fragments was produced at the University of Michigan Vector Core. Briefly, AAV helper vector pAd-Delta6 (200 µg), serotype vector pAAV2/8 (150 µg), and transfer pAAV-MCS2 vector carrying wild-type or Del52 knock-in HDR template (100 µg) were co-precipitated using standard PEI precipitation methods. PEI precipitation was performed by incubating plasmids with 270 µg PEI (molecular weight 2500, Polysciences, Inc.) in 60 mL Opti-MEM™ I reduced serum media (Life Technologies) at room temp for 15 min, before adding to fresh 60 mL DMEM, 10% FBS media. This DNA/PEI-containing media was then distributed equally to three T300 flasks (Falcon) containing HEK293T cells. Supernatant was collected at 72 h and frozen. Fresh DMEM with 5% FBS was then added, and both supernatant and cells were collected 48 h later. Cells were pelleted out and resuspended in 10 mL of Buffer A (150 mM NaCl2, 1 mM MgCl2, 50 mM Tris pH 8.5), frozen at -80°C for 20 min, thawed, and sonicated to release AAV particles. Following sonication, cell debris was pelleted out, and the supernatant was combined with the previously collected supernatant. AAV was then precipitated and pelleted out using PEG 8000 (Fisher). The pellet was resuspended in 10 mL Buffer A. The resuspended AAV was run through an iodixanol gradient (60%, 40%, 25%, and 15%). The 40% fraction was collected and applied to a 4 mL, 100kDa MWCO Amicon centrifugal filter and washed with PBS to remove iodixanol and concentrate the virus. Titers were assessed by qPCR using the protocol published by Addgene (https://www.addgene.org/protocols/aav-titration-qpcr-using-sybr-green-technology/).

**Generation of cell lines with heterozygous knock-in of the Del52 mutation**

Purified SpCas9-NLS was purchased from the QB3 MacroLab at the University of California, Berkeley. The Cas9 and sgRNA for CRISPR knock-in were complexed at a ratio of 2.5:1 (sgRNA: Cas9) at 37°C for 10 minutes to make ribonucleoprotein (RNP) complex. Cells were nucleofected with the RNP using Lonza 4D-nucleofector and the manufacturer's recommended protocol. THP-1 cells were nucleofected using the SG cell line kit and pulse code FF-100, and K562 cells were nucleofected using the SF cell line kit and FF-120 pulse code. Around 1 million cells were resuspended in 100 µL of nucleofector solution reconstituted with the provided supplement and mixed with the RNP complex containing 0.2 nmol of Cas9 and 0.5 nmol of sgRNA. Cells were nucleofected using the specific pulse code mentioned above and rested in 1 mL fresh complete media for 10 minutes. The cells were transduced with rAAV packaged with wild-type and mutant HDR templates at a multiplicity of infection of 10000:1 (vector genomes: cell). The virus-containing medium was replaced with fresh complete medium after 24 h. Cells were harvested, and the expression of reporter genes was detected by flow cytometry after 48 h. Populations of cells expressing GFP, BFP, or both, as well as those negative for the expression of both the reporter proteins, were sorted. Single cell clones of CRT-KO cells were grown by plating the cells in a 96-well plate at a density of one cell per two wells in the complete growth medium

**PCR-based confirmation of seamless integration of knock-in fragments at the endogenous *CALR* locus**

Genomic DNA was isolated from the single-cell clones of K562 or THP-1 cells sorted after CRISPR-based knock-in of wild-type *CALR* or $CRT_{Del52}$-specific exon 8,9 fragments. PCR was set up using genomic DNA as the template, a common forward primer that binds a *CALR* region upstream of the start site of the left homology arm and a specific reverse primer that binds to a sequence within the exon 9 of wild-type knock-in, $CRT_{Del52}$ knock-in, or endogenous *CALR* allele (supplementary table 3). The PCR amplicons were verified by agarose gel electrophoresis and Sanger Sequencing after gel purification.

**Generation of monoallelic HLA class I-expressing K562 cells**

The pMSCV neo constructs of HLA class I alleles were used for the expression of individual HLA class I allotypes in K562 cells. HEK293T cells were seeded in a six-well plate at a 0.5 million cells/well density and transfected with pMSCV-HLA construct (2.75 µg) along with 2 µg of pCL-Eco and 0.25 µg of VSV-g packaging and envelope plasmids in Opti-MEM™ I reduced serum media using FuGENE® HD (Promega) transfection reagent (10 µL). Virus-containing supernatant was collected after 48 h, filtered using 0.45 µm syringe filters to remove any HEK293T cells, and added to 0.5 million target K562 cells. Infection was done by spinoculation at 2500 rpm for 2h at room temperature in the presence of polybrene at 8 µg/mL final concentration. Half the volume of virus-containing media was replaced with the same volume of complete growth media after spinoculation, and the media was completely changed after 24 h of infection. The Geneticin™ (Gibco) was added 48 h after infection at the concentration mentioned above for the selection of infected cells. This procedure was also used for expression of HLA-B*44:02 and HLA-B*57:01 in K562 $GFP^-BFP^+$ cells with a heterozygous Del52 knock-in. For the expression of HLA-B*15:01, HLA-B*08:01, and HLA-B*07:02 in K562 $GFP^-BFP^+$ cells with heterozygous

Del52 knock-in, the pSIP constructs of the respective HLA class I genes were used. The HEK293T cells were transfected in a six-well plate with 2.25 µg pSIP-*HLA-B* vector along with 1.7 µg of psPAX2 and 1.1 µg of pMD2.G packaging and envelope plasmids. The Lipofectamine™ LTX reagent (15 µL) and Plus™ reagent (5 µL) were mixed with the plasmids in Opti-MEM™ I reduced serum media and incubated at room temperature for 10-15 minutes before adding the transfection complexes to the HEK293T cells. Various single-cell clones of K562 GFP$^-$ BFP$^+$ cells as well as the K562 parental cells were infected with the harvested virus in the same way as described above.

**CRISPR/Cas9-based CRT knockout and reconstitution of expression**

The HEK293T cells were transfected in a six-well plate with either 2.25 µg pLB empty vector or pLB carrying *CALR*-specific sgRNA (described in DNA constructs) along with 1.7 µg of psPAX2 and 1.1 µg of pMD2.G packaging and envelope plasmids. The Lipofectamine™ LTX reagent (15 µL) and Plus™ reagent (5 µL) were mixed with the plasmids in Opti-MEM™ I reduced serum media and incubated at room temperature for 10-15 minutes before adding the transfection complexes to the cells. Virus-containing supernatant was harvested 48 h after transfection, filtered to remove any HEK293T cells using 0.45 µm syringe filters, and added to 0.5 million target cells. Infection was done by spinoculation at 2500 rpm for 2h at room temperature in the presence of 8 µg/mL polybrene. The viral supernatant was replaced by complete media 24 h after infection, and cells were selected using blasticidin at the concentration mentioned above. Single-cell clones of CRT-KO cells were grown by plating the cells in a 96-well plate at a density of one cell per two wells in the complete growth medium.

For reconstitution of wild-type CRT or mutant CRT expression in CRT-KO cell lines, HEK293T cells were transfected with the psPAX2 and pMD2.G packaging and envelope plasmids along with 2.25 µg of sgRNA-resistant constructs of pSIP-$CRT_{WT}$, pSIP-$CRT_{Del52}$ or pSIP-$CRT_{Ins5}$, or the pSIP empty vector (see the DNA constructs section). The CRT-KO cells were infected with the harvested virus as explained above.

**qRT-PCR**

RNA was isolated from cells using the Qiagen RNeasy® Plus Mini kit (Catalog #74134) following the manufacturer's protocol. RNA was quantified by nanodrop, and the first strand of cDNA was synthesized from equal amounts of each RNA sample using the SuperScript™III First-Strand synthesis kit (Invitrogen; Catalog #18080051) with Oligo-dT primers, followed by digestion with RNase H enzyme for 20 minutes at 37°C. The cDNA was either used immediately or stored at -80°C. For qPCR, 1 µL of cDNA was mixed with the SYBR™ Green PCR master mix (1X) (Applied Biosystems, catalog #4309155) and 500 nM each of forward and reverse primers targeted against the wild-type *CALR* exon 9 (supplementary table 3) or the housekeeping genes. The qPCR was performed in an Applied Biosystems 7500 Fast Real-Time PCR machine. The primers for human GAPDH (catalog #VHPS-3541), human beta actin (catalog #VHPS-4263), and human hypoxanthine phosphoribosyltransferase 1 (HPRT1) (catalog #VHPS-3541) housekeeping genes were procured from Real Time Primers, LLC.

**PBMC and platelet isolation**

Whole blood samples from de-identified patients or healthy donors were received in BD Vacutainer® ACD solution A tubes (catalog number: 364606). PBMC were isolated from whole blood samples by density gradient centrifugation on Ficoll® Paque-Plus (Cytiva-Millipore Sigma) media. Blood sample was diluted to 35 mL with PBS+2% FBS and layered slowly on 15 mL Ficoll® Paque-Plus density gradient media and centrifuged at 400g for 30 minutes at room temperature using 4 acceleration and 0 deceleration settings. The buffy layer with PBMC was collected and washed thrice with PBS + 2%FBS. PBMC were resuspended in complete RPMI media, counted, and used on the same day as the blood collection for staining of HLA class I or surface CRT for flow cytometry, as explained below, or frozen in the Recovery™ Cell Culture Freezing Medium (Gibco) for other experiments.

Platelets were isolated from whole blood before PBMC isolation as described previously [9,10]. Platelet-rich plasma (PRP) was prepared from the whole blood sample by centrifugation on a swing bucket rotor at slow speed (200 x g) for 15 min at room temperature (RT) using 2 acceleration and zero deceleration settings. The yellow-colored layer of PRP on the top of the red blood layer was carefully transferred to a fresh polypropylene tube using a transfer pipette without disturbing the layer of blood at the bottom. The PRP was treated with 0.02 U/mL apyrase and ACD solution (Sigma, 1:10 dilution). The PRP was centrifuged at 2000 x g for 10 min at RT at slow (~4) acceleration/deceleration settings to separate the platelets from cleared plasma. Platelets were rested in ~10 mL of Tyrode's buffer (10 mM HEPES pH 7.4, 134 mM NaCl, 12 mM $NaHCO_3$, 2.9 mM KCl, 0.34 mM $Na_2HPO_4$, 1 mM $MgCl_2$ and 5 mM D-glucose) for 30 min at 37°C before being used for surface HLA class I measurements or making lysates for immunoblotting.

**HLA genotyping**

Genomic DNA was isolated from donor PBMC using the DNeasy® Blood and Tissue Kit from Qiagen (Catalog #69504) following the manufacturer's protocol. For HLA genotyping, genomic DNA samples were submitted to Sirona Genomics (Immucor, USA). The MIA FORA NGS MFLEX XP HLA Typing Kit (Werfen, USA) was used that utilizes long-range PCR to amplify all exons of clinically significant Class I (*HLA-A, -B, -C*) and Class II (*HLA-DRB, -DQB1, -DPB1, -DPA1, -DQA1*) HLA genes in a multiplex reaction for high-resolution HLA typing via NGS sequencing. The resulting PCR products were prepared for sequencing with the MIA FORA Library Kit. The sequencing was performed on the Illumina MiniSeq platform. The generated sequences were then analyzed using MIA FORA NGS Software v5.3.0 with database 3500 to determine HLA typing results.

**Antibodies**

The antibodies used in this study have been listed below, along with the source, dilution, and the specific experiment for which each of them was used.

| S.No | Antibody description | Vendor | Catalog number | Dilution | Experiment |
|---|---|---|---|---|---|
| 1 | Anti-CD3 Pacific Blue | BioLegend | 300431 | 1:200 | Surface staining of PBMC |
| 2 | Anti-CD19 APC | BioLegend | 302212 | 1:200 | Surface staining of PBMC |
| 3 | Anti-HLA-DR BV650 | BioLegend | 307650 | 1:200 | Surface staining of PBMC |
| 4 | Anti-CD14 AlexaFluor®700 | BioLegend | 367114 | 1:200 | Surface staining of PBMC |
| 5 | Anti-CD16 PE | BioLegend | 302008 | 1:200 | Surface staining of PBMC |
| 6 | Anti-CD41 AlexaFluor® 647 | BioLegend | 303726 | 1:50 | Surface staining of platelets |
| 7 | W6/32-FITC | Purified in lab from mouse ascites and conjugated to FITC | | | Surface staining of PBMC, platelets and cell lines |
| 8 | W6/32-PE | BioLegend | 311406 | 1:100 | Surface staining of cell lines |

| | | | | | |
|---|---|---|---|---|---|
| 9 | Anti-HLA-A2 (BB7.2) PE/Cy7 | Biolegend | 343314 | 1:200 | Surface staining of PBMC, platelets and cell lines |
| 10 | Anti-Bw4 FITC | One Lambda | FH0007 | 1:20 | Surface staining of cell lines |
| 11 | Anti-Bw6 FITC | One Lambda | FH0038 | 1:50 | Surface staining of cell lines |
| 12 | Anti-CRT(N) | Cell Signaling Technologies | 12238S | 1:100 Surface CRT; 1:10,000 Immunoblots | Immunoblots and surface CRT staining for flow cytometry |
| 13 | Anti-CRT($C_{mut}$) | Custom-generated[5] | | 1:100 Surface CRT; 1:10,000 Immunoblots | Immunoblots and surface CRT staining for flow cytometry |
| 14 | Anti-GAPDH | Cell Signaling Technologies | 2118S | 1:10000 | Immunoblots |
| 15 | Anti-Vinculin | Cell Signaling Technologies | 13901S | 1:10000 | Immunoblots |
| 16 | FITC Mouse IgG2a Isotype control antibody | BioLegend | 400208 | 1:200 | Surface staining of PBMC, platelets and cell lines |
| 17 | PE-Mouse IgG2a Isotype control antibody | BioLegend | 400214 | 1:100 | Surface staining of cell lines |
| 18 | FITC-Mouse IgG3 Isotype control antibody | Abcam | ab91539 | 1:50 | Surface staining of cell lines |
| 19 | PE/Cy7-Mouse IgG2b Isotype control antibody | BioLegend | 400326 | 1:200 | Surface staining of PBMC, platelets and cell lines |

**Cell lysis and immunoblotting**

Cells were collected and washed with 1X PBS (room temperature). The pelleted cells were lysed in 100-200 µL lysis buffer (50 mM Tris pH 7.5, 150 mM NaCl, 1% Triton X-100, 5 mM $CaCl_2$ supplemented with protease inhibitor cocktail (P8340 from Sigma) at 1:100 dilution) for 30 min at 4°C. BCA assay (Pierce) was used to quantify the total protein in the lysates. The lysates were boiled with 1X SDS loading dye for 5 minutes at 99°C. Platelets lysates were either prepared on the day of blood collection, or the platelet pellets were stored at -80°C and lysed later using the same lysis buffer mentioned above. The lysates were quantified for total protein and boiled with 1X SDS loading dye and stored at -80°C until use. The lysates were subjected to SDS-PAGE and western blotting. Wild-type and mutant CRT proteins were detected using anti-CRT(N) and anti-CRT($C_{mut}$) antibodies, respectively. Vinculin or GAPDH protein levels in the lysates were probed loading control. Blots were incubated with Horseradish peroxidase-conjugated secondary antibodies (Jackson ImmunoResearch) and developed via chemiluminescence (Pierce™ ECL western blotting substrate, 32106). Image J was used for the quantification of band intensities on Western blots.

**Flow cytometry and antibody binding capacity (ABC) measurements**

The protocol for surface HLA class I staining on PBMC has been described previously[11]. Freshly isolated PBMC were stained with antibodies against specific surface markers and surface HLA class I for 30 minutes at 4°C in FACS buffer (PBS + 2% FBS). The antibodies (details in the antibodies section) against surface markers included anti-CD3 Pacific Blue, anti-CD19 APC, anti-HLA-DR BV650, anti-CD14 AlexaFluor™ 700, and anti-CD16 PE. Platelets were stained with anti-CD41 AlexaFluor™ 647 and the specific anti-HLA class I antibody or the corresponding isotype control for 30 min at room temperature in FACS buffer. Surface HLA class I staining was performed with the W6/32-FITC antibody, the PE/Cy7-conjugated anti-HLA-A2 (BB7.2) antibody, or the corresponding isotype control antibodies. For surface CRT staining, PBMC were stained with anti-CRT(N) antibody or anti-CRT($C_{mut}$) antibody in FACS buffer for 30 min at 4°C, followed by staining with AlexaFluor™ 647-conjugated goat anti-rabbit secondary antibody (1:500; Invitrogen) for 15 minutes at 4°C. Dead cells were stained with 7-Amino-Actinomycin D (7-AAD) dye (BD, catalog #559925). For quantitative flow cytometry experiments, Quantum™ simply cellular anti-mouse IgG beads (Bangs Laboratory, Inc.) were stained concurrently with the cells using the same antibody dilution. Standard curves were plotted using the geometric mean fluorescence intensity (gMFI) values for various bead populations *vs.* the number of Fc receptors on the respective beads provided by the manufacturer. The antibody binding capacity (ABC) values for different cell types were interpolated from the standard curve.

The cell lines were stained with the W6/32, anti-HLA-A2 (BB7.2), anti-Bw6 or anti-Bw4 antibody for 30 min at 4°C in FACS buffer. The cells were washed twice with FACS buffer followed by staining with 7-AAD viability dye. THP-1 cells were blocked with 5% Normal Mouse Serum (Jackson laboratories) in FACS buffer for 10 minutes at room temperature before surface staining with the anti-HLA class I antibodies as given above. Cells were analyzed using either BD LSRFortessa™ or BD FACSCanto™ cell analyzer. The analysis of flow cytometry data was performed in FlowJo™ version 10.

**Variant allele frequency analysis**

The wild-type or mutant *CALR*-specific exon 9 fragments were amplified using the primers given in the supplementary table 3, and genomic DNA isolated from MPN patient PBMC as template. The amplified fragments were purified by Qiagen PCR purification kit (Catalog #28106). The purified amplicons were submitted to Genewiz (Azenta Life Sciences) for variant allele frequency analysis using the Amplicon-EZ next-generation sequencing service.

**FlowPRA screening of BB7.2 binding specificities**

The HLA class I-binding specificities of the BB7.2 antibody were checked by using the FlowPRA single antigen HLA class I antibody detection test (One Lambda). We checked for binding to groups of beads coated with antigens derived from common HLA-A, B, and C alleles, including FlowPRA single antigen beads Group 1,2,3,4,6,9, and 10. The beads were vortexed, and 5 µL of beads were incubated with 20 µL of the BB7.2-PE/Cy7 antibody at a 1:200 dilution for 30 minutes at room temperature, followed by two washes with the 1X dilution of wash buffer (included in the kit) in water. The beads were resuspended in 1X PBS and analyzed by flow cytometry. Each group contains 9 bead populations that exhibit different levels of fluorescence in the PE channel. Additionally, each group includes negative control beads without antigens that can be used to set the threshold for negative binding.

**References**

1. Foßelteder J, Pabst G, Sconocchia T, et al. Human gene-engineered calreticulin mutant stem cells recapitulate MPN hallmarks and identify targetable vulnerabilities. *Leukemia*. 2023;37(4):843-853.
2. Mohan HM, Yang B, Dean NA, Raghavan M. Calreticulin enhances the secretory trafficking of a misfolded alpha-1-antitrypsin. *J Biol Chem*. 2020;295(49):16754-16772.
3. Fosselteder J, Pabst G, Sconocchia T, et al. Human gene-engineered calreticulin mutant stem cells recapitulate MPN hallmarks and identify targetable vulnerabilities. *Leukemia*. 2023;37(4):843-853.
4. Klampfl T, Gisslinger H, Harutyunyan AS, et al. Somatic Mutations of Calreticulin in Myeloproliferative Neoplasms. *N Engl J Med*. 2013;369(25):2379-2390.
5. Venkatesan A, Geng J, Kandarpa M, et al. Mechanism of mutant calreticulin-mediated activation of the thrombopoietin receptor in cancers. *J Cell Biol*. 2021;220(7):e202009179.
6. Bashirova AA, Viard M, Naranbhai V, et al. HLA tapasin independence: broader peptide repertoire and HIV control. *Proc Natl Acad Sci U S A*. 2020;117(45):28232-28238.
7. Garcia-Beltran WF, Holzemer A, Martrus G, et al. Open conformers of HLA-F are high-affinity ligands of the activating NK-cell receptor KIR3DS1. *Nat Immunol*. 2016;17(9):1067-1074.
8. Rizvi SM, Salam N, Geng J, et al. Distinct Assembly Profiles of HLA-B Molecules. *The Journal of Immunology*. 2014;192(11):4967-4976.
9. Greening DW, Sparrow RL, Simpson RJ. Preparation of platelet concentrates. *Methods Mol Biol*. 2011;728:267-278.
10. Kaur A, Venkatesan A, Kandarpa M, Talpaz M, Raghavan M. Lysosomal degradation targets mutant calreticulin and the thrombopoietin receptor in myeloproliferative neoplasms. *Blood Adv*. 2024;8(13):3372-3387.
11. Yarzabek B, Zaitouna AJ, Olson E, et al. Variations in HLA-B cell surface expression, half-life and extracellular antigen receptivity. *Elife*. 2018;7.
